# From defaults to databases: parameter and database choice dramatically impact the performance of metagenomic taxonomic classification tools

**DOI:** 10.1101/2022.04.27.489753

**Authors:** Robyn J. Wright, André M. Comeau, Morgan G.I. Langille

**Affiliations:** Department of Pharmacology, Faculty of Medicine, Dalhousie University, Halifax, Canada; Integrated Microbiome Resource (IMR), Dalhousie University, Halifax, Canada

## Abstract

In metagenomic analyses of microbiomes, one of the first steps is usually the taxonomic classification of reads by comparison to a database of previously taxonomically classified genomes. While different studies comparing metagenomic taxonomic classification methods have determined that different tools are “best”, there are two tools that have been used the most to-date: Kraken (*k*-mer based classification against a user-constructed database) and MetaPhlAn (classification by alignment to clade-specific marker genes), the latest versions of which are Kraken2 and MetaPhlAn 3, respectively. We found large discrepancies in both the proportion of reads that were classified as well as the number of species that were identified when we used both Kraken2 and MetaPhlAn 3 to classify reads within metagenomes from human-associated or environmental datasets. We then investigated which of these tools would give classifications closest to the real composition of metagenomic samples using a range of simulated and mock samples and examined the combined impact of tool-parameter-database choice on the taxonomic classifications given. This revealed that there may not be a one-size-fits-all “best” choice. While Kraken2 can achieve better overall performance, with higher precision, recall and F1 scores, as well as alpha- and beta-diversity measures closer to the known composition than MetaPhlAn 3, the computational resources required for this may be prohibitive for many researchers, and the default database and parameters should not be used. We therefore conclude that the best tool-parameter-database choice for a particular application depends on the scientific question of interest, which performance metric is most important for this question and the limit of available computational resources.

## 1. Introduction

One of the first steps in most analyses of microbiomes characterised using metagenomic sequencing (MGS) is to taxonomically classify the reads present in each sample. There are many metagenomic classification programs that have been developed to do this, and while these use a range of different methods, one common step is the comparison of unknown reads within a sample to a database of known genomes or sequences. Several studies have compared the performance of different metagenomic classifiers using simulated or mock communities with a known taxonomic composition [1–4] or a consensus approach [5, 6] and publications introducing new metagenomic classifiers also often compare the method being introduced with previous methods [7–9]. The “best” metagenomic classifier is frequently determined based on either the F1 score (harmonic mean of precision and recall) or L1 distance (also known as Manhattan or Taxicab distance), and typically varies depending on the environment that a sample comes from as well as the complexity (number of taxa and magnitude of abundance differences) of the sample [1, 8]. Different comparison studies have therefore concluded that different metagenomic classifiers are “best”.

Another factor that can have a large impact on the taxonomic classifications obtained from any given metagenomic classifier is the choice of database used for comparison. It is largely established that the larger and more comprehensive the database used for comparison, the better, to avoid erroneous classifications caused by the omission of close relatives for a given taxon [10], but this represents a significant computational burden. For example, in version 205 (May 2021) of the NCBI RefSeq database [11] there were 108,257 organisms (227,889 genomes), including 64,833 bacterial species. This full database uses approximately 1.2 TB of disk space for storage and has similar memory requirements, leading many researchers to use a reduced version of this database. However, to our knowledge, there are few microbial metagenome studies that compare more than one database [1, 3, 8], and we were unable to find a systematic characterisation of the effect that this has on classification accuracy.

Regardless of the metagenomic classification program that performs the “best”, the two programs that have been used the most to date (based on number of citations per year that they have received; Fig. S1) are Kraken [12] and MetaPhlAn [13]. MetaPhlAn (<u>Meta</u>genomic <u>Ph</u>ylogenetic <u>An</u>alysis) was first introduced in 2012 [13] and the current version, MetaPhlAn 3 (introduced in 2021 [7]), uses a database of clade-specific marker genes from 13,475 taxa to assign query reads to clades using Bowtie 2 [14]. Kraken was first introduced in 2014 [12] and the current version, Kraken2 (introduced in 2019 [15]), uses exact alignment of *k*-mers (sub-sequences of length *k*) to a reference database for query read classification. Because *k*-mers within a read could map to multiple sequences within the database when using Kraken2, a lowest common ancestor (LCA) approach is used to classify the sequence, and the number of *k*-mers mapping to a given taxon are calculated as a proportion of the number of *k*-mers within that sequence that do not have ambiguous nucleotides. This proportion of *k*-mers mapping to a taxon is called the “confidence”, and there is a confidence threshold that can be set within the program (0 by default), where a taxonomic classification is only taken if it is above this pre-defined threshold. Furthermore, the Kraken creators also suggest using Bracken (<u>B</u>ayesian <u>R</u>eestimation of <u>A</u>bundance after <u>C</u>lassification with <u>K</u>rak<u>en</u>) [16], which combines the classifications made by Kraken with information about the genomes within the database to estimate the abundance of taxa within a sample.

The recommendations to set a confidence threshold and to use Bracken following Kraken are not given in either the original [12] or Kraken2 [15] publications. We knew anecdotally that the confidence threshold had a large impact on the number of reads classified as well as the number of taxa found to be present within a sample, but there is not, to our knowledge, a published study that investigated it systematically. The confidence threshold is discussed in the online manual for Kraken2 (although guidelines on choosing an appropriate threshold are limited) while Bracken is introduced in a separate publication [16]. We believe that this is the reason that even studies comparing metagenomic classifiers – that use Kraken2 – don’t always follow the recommendation to use Bracken, *e*.*g*., [2]. Furthermore, we only found three studies that mentioned setting a confidence threshold within Kraken2, all comparisons of metagenomic classifiers that set confidence thresholds of 0.05 [3] (Kraken2), 0.2 [1] (Kraken1) or 0.5 [17] (Kraken2), although if these thresholds were investigated further, this was not presented in the publications. Also, the creators of Kraken2 provide a reduced database alongside instructions for building a database from all genomes contained in the NCBI RefSeq database, but many studies use some form of a reduced database [1, 3], or we were unable to find details of the database used at all [5, 9, 17, 18], or with enough detail to replicate [2, 4].

Here we initially compared the results given by both Kraken2 (with Bracken) and MetaPhlAn 3 on real MGS datasets that had paired 16S rRNA gene (16S) data available. Due to the large discrepancies between the methods, we used a range of simulated and mock MGS samples to systematically investigate the impact of database and confidence threshold choice on Kraken2 performance. Finally, after determining the optimal parameters for running each tool, we compared the classifications given by both Kraken2 and MetaPhlAn 3 on the simulated and mock samples and we give guidance on the scenarios in which we think each tool is most appropriate.

## 2. Methods

### 2.1. Datasets used and sample pre-processing

#### 2.1.1. Real 16S and MGS dataset characteristics

We used datasets (Table S1A) that had both metagenome and 16S rRNA gene sequencing data available, as used previously in Douglas *et al*. [19] (details of accession numbers or where to download data are given in Table S1A):

1. Cameroonian [20] (*n*=57): stool samples from Cameroonian individuals.
2. HMP [21] (*n*=137): the Human Microbiome Project healthy samples.
3. Indian [22] (*n*=91): stool samples from Indian individuals.
4. Mammal [23] (*n*=11): stool samples from mammals.
5. Ocean [24] (*n*=2): ocean water samples.
6. Primate [25] (*n*=74): non-human primate stool samples.
7. Soil [26] (blueberry; *n*=14): bulk soil and blueberry rhizosphere samples.

The processed 16S files for all datasets and processed MGS files for the Cameroonian, HMP, Indian and Soil (blueberry) datasets as well as raw MGS files for the Mammal, Ocean and Primate datasets were provided by Douglas *et al*. [19]. These are split to human-associated (datasets 1, 2 and 3) and environmental (datasets 4, 5, 6 and 7) in some analyses.

#### 2.1.2. Simulated and mock sample dataset characteristics

We used several datasets (Table S1B) containing simulated or mock samples (*n*=389) that have been generated by previous metagenomic classifier benchmarking studies (details of accession numbers and where files were downloaded from are given in Table S1B):

1. Simulated (*n*=26 with 0-525 taxa per sample), mock biological (*n*=26 with 0-23 taxa per sample) and mock biological/simulated (*n*=3, 113 taxa per sample) samples used by McIntyre *et al*. [1] where the samples were selected from previous studies [27–32]. For each of these, the reads were either simulated, generated from samples where the composition was known prior to DNA extraction or sequences or contigs were assembled from sequenced isolates, respectively.
2. Simulated samples made by Parks *et al*. [8] to have: (i) an average of either 100 (low) or 500 (high) species per sample; (ii) one (low) or up to 10 (high) strains per species; (iii) average nucleotide identity (ANI) to reference genomes of 100% (identical), 99% (high), 97% (moderate) or 95% (low). They constructed ten replicates for each of these aside from 95% ANI with high species diversity, totalling 140 samples.
3. The low (*n*=1), medium (*n*=2, with short circular element insert sizes) and high (*n*=5) complexity datasets generated for the Critical Assessment of Metagenome Interpretation (CAMI) [5].
4. Short-read samples simulated for the marine (*n*=10), rhizosphere/plant-associated (*n*=21) or mouse gut (*n*=49) environments as well as the strain madness (*n*=100) dataset from the second Critical Assessment of Metagenome Interpretation challenges (CAMI2) [6].

We also used one ZymoBIOMICS Microbial Community Standard (Zymo Research, USA, containing three gram-positive and five gram-negative bacteria and two yeasts). The DNA from this sample was extracted and sequenced on an Illumina NextSeq by the Integrated Microbiome Resource (IMR, Dalhousie University) following the standard operating procedures in Comeau *et al*. [33]. Samples from datasets 1 and 2 had known compositions in terms of the number of reads that were from different genomes (simulated samples), or the proportion of the sample that was composed of different species (mock samples; either before or after DNA extraction). The CAMI and CAMI2 samples (datasets 3 and 4) had a “gold standard” taxonomic profile, calculated by the original authors, as well as information on the provenance of individual reads for CAMI2 only (dataset 4). Briefly, they created the gold standard taxonomic profiles by: (a) extracting the marker genes from the genomes of sequenced isolates (that were subsequently used for sample construction), (b) alignment of the 16S rRNA marker genes to a reference collection, (c) alignment and clustering of these sequences, and (d) taxonomic annotation and Operational Taxonomic Unit (OTU; 3% distance threshold) assignment based on the clustering of marker genes. Each genome is also assigned to a novelty category based on how closely related it is to other genomes in the reference database (*i*.*e*., whether it represents a new strain, species, etc.). The gold standard taxonomic profiles are given as relative abundances at each taxonomic rank and samples may therefore not sum to 100% at every rank due to the presence of novel genomes.

#### 2.1.3. Sample processing

Where necessary, Kneaddata [7] (version 0.7.4) was run on samples for quality control and primer trimming using the --very-sensitive --dovetail Bowtie 2 [14] options and paired-end reads were combined to a single file prior to further analysis.

### 2.2. Database construction

#### 2.2.1. MetaPhlAn 3

The MetaPhlAn 3 database used was the default database, downloaded using the install commands within MetaPhlAn 3. This database is based on the ChocoPhlAn 3 database, which contains approximately 99,200 fully annotated genomes that span 561 archaeal, 9,795 bacterial, 122 eukaryotic, and 2,997 viroid/viral taxa [7]. The species-level pangenomes are then used to identify core gene families that are present in almost all strains of a species and these markers are what is used in the MetaPhlAn 3 database (2.8 GB).

#### 2.2.2. Kraken2

Kraken2 provides commands for the construction of default databases using the NCBI RefSeq database (and will by default use the most recent release). Their standard database (described in the Kraken2 publication [15]) includes complete genomes from the bacteria, archaea, viruses, the human genome and a collection of known vectors (UniVec_Core). The only databases provided by the Kraken2 team in a ready-to-use format are the MiniKraken V1 (without the human genome) and V2 (with the human genome) databases, which are versions of the standard database where minimizers are sub-sampled to reduce the database size to 8 GB. We wanted to investigate the effect that the database used has on the performance of taxonomic classification in a systematic manner. We therefore based our Kraken2 databases on the NCBI RefSeq database (release 205, May 2021), containing 108,257 organisms and 227,854 genomes for 1,131 Archaeal, 214,594 Bacterial, 1,266 Eukaryotic and 10,863 Viral genomes, and on the Genome Taxonomy Database (GTDB; release 202, April 2021), containing 2,339 Archaeal and 45,555 Bacterial species. We also used the nt (non-redundant nucleotide sequences), Plasmid and UniVec_Core sequences. We initially downloaded all genomes and used custom scripts written in Python to filter out the genomes necessary to build each database and then used the Kraken2 build commands for construction of the databases (final disk sizes of each database are given in parentheses):

1. NCBI RefSeq Complete V205 (1,189 GB): all 227,889 genomes included in release 205 of the NCBI RefSeq as well as the nt, Plasmid and UniVec_Core sequences.
2. NCBI RefSeq Complete V205 500 GB (466 GB): constructed as for NCBI RefSeq Complete V205 but with the --max-db-size command to create a database with a maximum size of 500 GB by sub-sampling minimizers.
3. NCBI RefSeq Complete V205 100 GB (94 GB): constructed as for NCBI RefSeq Complete V205 but with the --max-db-size command to create a database with a maximum size of 100 GB by sub-sampling minimizers.
4. NCBI RefSeq V208 non-redundant nucleotide (nt) database (308 GB): we were unable to use V205 for this database as it was constructed at a later date than the other databases and previous versions are unavailable on the NCBI FTP site. The number of organisms included had increased from 108,257 for V205 to 113,002 for V208. This was created using only Kraken2 commands and not the genomes described above.
5. GTDB r202 bacteria/archaea + NCBI RefSeq V205 other domains (1,148 GB): all bacterial and archaeal genomes (*n*=60,068) included in release 202 of GTDB as well as the genomes for other domains included in release 205 of NCBI RefSeq (*n*=12,176).
6. ChocoPhlAn 3 (73 GB): all genomes (*n*=132,661) matching taxon IDs (*n*=13,475) included in the ChocoPhlAn 3 database used for MetaPhlAn 3.
7. Standard (51 GB): the standard database provided in a ready to use format and linked from the Kraken2 website (https://ccb.jhu.edu/software/kraken2/index.shtml) that is available from the Langmead lab website (https://benlangmead.github.io/aws-indexes/k2; 17 May 2021 version) and contains complete genomes from 377 Archaea, 21,543 Bacteria and 10,489 Viruses as well as the Plasmid and UniVec_Core sequences and the human genome.
8. Minikraken2 V2 (8 GB): a reduced database available from the Kraken2 website (https://ccb.jhu.edu/software/kraken2/index.shtml) that includes 159 Archaeal, 4,107 Bacterial and 1,491 Viral taxa and the human genome. Note that the number of genomes that this corresponds to is not given by the Kraken2 team and the taxa included were determined using the Kraken2 database-inspect command. We were not able to find the release of NCBI RefSeq from which this was built.

The accession number and taxonomy ID for all genomes included as well as which databases each genome was used for are given in Table S2.

### 2.3. Taxonomic classification of samples

#### 2.3.1. Taxonomic classification of real 16S and MGS dataset

The MGS validation datasets were run using Kraken2 followed by Bracken at the species and genus levels with each of the described databases with confidence thresholds of 0.00 or 1.00 as well as with MetaPhlAn 3 with the -t rel_ab_w_read_stats option. We used the scikit-learn module within QIIME2 v2021.11 [34] with a classifier trained on the full-length 16S rRNA gene Silva V138 database [35–37] to classify the 16S rRNA gene sequence files.

#### 2.3.2. Kraken2 classification of simulated and mock samples

Kraken2 [15] was run on all simulated and mock samples with each of the described databases using the default commands and a confidence threshold of between 0.00 and 1.00 at intervals of 0.05. After running Kraken2, Bracken [16] was run at the species level to re-estimate the abundance of taxa. Additionally, Kraken2 (followed by Bracken) was also run with the experimental --report-minimizer-data feature, adapted from KrakenUniq [38], with a confidence threshold of 0.00 and the RefSeq Complete V205 database only. The number of distinct minimizers per taxon in the Kraken2 output was used to filter the Bracken output by keeping only taxa with more than 1, 5, 10, 100, 500, 1,000, 2,500, 5,000, 10,000, 25,000, 50,000 and 100,000 distinct minimizers.

#### 2.3.3. MetaPhlAn 3 / HUMAnN 3 classification of simulated and mock samples

MetaPhlAn 3 [7] was run in one of several ways: (**1**) using the default parameters and multiplying the resulting relative abundances by the number of reads in each sample; (**2**) using the default parameters with the -t rel_ab_w_read_stats option for estimating the number of reads from each clade; (**3**) using different options for Bowtie2 mapping of reads to the ChocoPhlAn 3 marker gene database with --bt2-ps, *i*.*e*., the default very-sensitive as well as the other options sensitive, sensitive-local and very-sensitive-local; (**4**) using the different statistical options for converting marker gene abundances into clade abundances with --stat, *i*.*e*., with the default truncated clade global average (tavg_g) and the other options of clade global average (avg_g), average of length-normalized marker counts (avg_l), truncated average of length-normalized marker counts (tavg_l), winsorized clade global average (wavg_g), winsorized average of length-normalized marker counts (wavg_l) and the median of length-normalized marker counts (med); or (**5**) by running HUMAnN 3 and examining the reads mapped with Bowtie2 in the nucleotide alignment step, *i*.*e*., reads mapped to the full genomes of the taxa identified by MetaPhlAn 3 or the reads mapped with Diamond [39] to the UniRef90 or UniRef50 [40] databases in the translated alignment step.

The additional programs GNU Parallel [41] and GNU Time were used to wrap commands and allowed multiple samples to be classified at the same time and the time taken for each sample to be classified to be recorded, respectively. A server running Ubuntu 16.04 with 1.5 TB RAM and 4x Intel(R) Xeon(R) CPU E7-8870 v4 @ 2.10GHz = 40 cores (80 threads) was used to build all databases and process all samples.

### 2.4. Statistical analyses on real datasets

The classifications obtained from either Kraken2 or MetaPhlAn 3 were combined at both the species and the genus levels for MGS samples and the proportion of reads that were classified as well as the proportion of reads that were classified as bacteria were taken from these outputs. For 16S samples, we took OTUs/ASVs as being approximately equivalent to the MGS species (only 5,169 of 21,453 16S sequences have a species-level classification) and used the genus-level classifications. For both 16S and MGS samples, the number of species and the number of genera that were present within each sample was calculated from the classifications.

### 2.5. Statistical analyses on simulated and mock samples

#### 2.5.1. Samples included

For the plotting and analyses shown here, we removed samples where the reads were too long *i*.*e*., from Nanopore sequencing (*n*=14) or were from assembled contigs (*n*=3), samples with reads that were simulated to map to multiple domains (*n*=2), samples simulated as negative controls (*n*=3) and one other sample with only 112 reads. The 11 samples that had known proportions of organisms or DNA as well as the CAMI and CAMI2 samples were analysed separately (see below), leaving 164 samples with known compositions that included the number of reads that should map to each taxon. These 164 samples with known numbers of reads were used for all analyses unless otherwise specified. We used the 11 samples with known proportions of organisms or DNA to determine how well each classification program was able to reconstruct this. For Kraken2 classifications, we normalised the reads output using the expected genome size for each taxon ID; NCBI gives an expected genome size for each species with at least four assemblies, so we obtained these estimates for all taxon IDs. For other taxon IDs, we took the median genome size of the genomes with this taxonomy ID. There were taxon IDs (2,793 of 25,126, accounting for a median of 0.004% of reads classified within a sample) for which we couldn’t get a genome size; in these cases, we used the median size of all genomes for normalisation. MetaPhlAn 3 already gives estimated relative abundance within the community as an output, so no further normalisation was carried out.

#### 2.5.2. Statistical analyses

First, the results for each sample were combined for each database or tool-parameter comparison and only the NCBI taxonomy ID was kept for each taxon. For the GTDB classifications, we converted the taxonomy ID structure that we had built according to the GTDB taxonomic classifications for genomes to NCBI taxonomy, although we note that some levels of the GTDB taxonomic structure do not have an NCBI equivalent (*e*.*g*., some genomes only have a strain and then a family assignment in the NCBI taxonomy, but have a genus and species assignment in the GTDB taxonomy). Next, taxonomic trees were constructed using PhyloT (https://phylot.biobyte.de), which incorporate phylogenetic and taxonomic knowledge, but not branch lengths or clade support values. In making the list of taxonomy IDs needed for the tree, taxonomy IDs that are no longer present in the NCBI taxonomy structure were either merged (*n*=65,504) or removed (*n*=2,102). For a small number of taxon IDs (*n*=1,046, 0.14% of reads) assigned using RefSeq V208, these taxa were instead assigned their parents taxon ID because these are not yet present in PhyloT (as of 10 November 2021). This left 214,504 taxonomy IDs. For each result, several metrics for assessing their accuracy were calculated:

1. ***Basic metrics***: 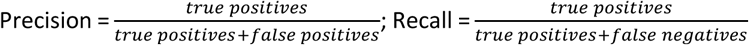 and 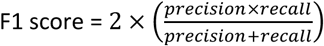
2. ***Alpha-diversity metrics***: Proportion of reads within the sample that were classified; Number of species that were identified; Simpson’s diversity [42]; Shannon diversity [43]; Faith’s Phylogenetic diversity [44]; Chao1 richness [45]; McIntosh’s evenness [46]; Pielou’s evenness [47]; and Simpson’s evenness [42].
3. ***Beta-diversity metrics***: L1 distance; Aitchison’s distance [48]; Robust Aitchison’s distance [49]; Bray-Curtis dissimilarity [50]; and Weighted and unweighted UniFrac distance [51].

In each case, the result given by a tool was compared with the known composition for each sample. For precision, recall and F1 score, values were calculated both in terms of the number of taxa and the number of reads, with the mean precision, recall and F1 scores calculated as the mean of the precision, recall and F1 scores, respectively, for reads and taxa. For Bray-Curtis dissimilarity and weighted and unweighted UniFrac distances, values were calculated both for unnormalized reads and relative abundance. Note that we refer to the Faith’s phylogenetic diversity and weighted and unweighted UniFrac distance as Faith’s taxonomic diversity and weighted and unweighted taxonomic UniFrac distance, respectively, to make clear that these metrics were calculated with the PhyloT taxonomic tree and not with a true phylogenetic tree. For alpha-diversity, richness and evenness metrics, we subtracted the true value of the sample from that calculated from the given classifications.

#### 2.5.3. CAMI sample analysis

Due to the samples that were constructed by both the CAMI (*n*=10) [5] and CAMI2 (*n*=180) [6] studies using newly sequenced genomes, not all genomes – and therefore not all reads – have classifications at all taxonomic ranks. For some genomes, the lowest rank of the taxonomic classification given is at the family or genus level. This therefore makes the calculation of many of the above metrics misleading, and we have consequently analysed these samples separately from the other simulated samples. For these samples we therefore examine only the proportion of reads classified (as compared with the gold standard taxonomic profile given, described above) and the recall for the number of taxa and for the number of reads. For the CAMI2 samples, in addition to the community-level relative abundance of taxa given as the gold standard, the taxonomic provenance is given on a read-by-read basis. As Kraken2 also includes an output where taxonomic assignments are given for each read, this allowed us to assign an index to each read based on whether the Kraken2 classification was:

i. The same as the CAMI2 gold standard taxonomic classification (index=0);
ii. At a lower rank than the CAMI2 gold standard taxonomic classification, but was the same at the rank of the CAMI2 gold standard taxonomic classification, *e*.*g*., Kraken2 assigned a read to the species *Ruminococcus albus*, but the CAMI2 gold standard taxonomic classification was the genus *Ruminococcus* (index=0);
iii. At a higher rank than the CAMI2 gold standard taxonomic classification, but this was a rank that the CAMI2 gold standard taxonomic classification belonged to, *e*.*g*., Kraken2 assigned a read to the *Saprospira* genus, but the CAMI2 gold standard classification was the species *Saprospira grandis* (index=1, as the Kraken2 classification is one rank away from the CAMI2 gold standard);
iv. Incorrect at the rank of the CAMI2 gold standard taxonomic classification, but both classifications shared a higher rank, *e*.*g*., Kraken2 assigned a read to the species *Corticicoccus populi*, but the CAMI2 gold standard taxonomic classification was the species *Staphylococcus aureus*, both of which belong to the *Staphylococcaceae* family (index=2, as the Kraken2 classification is correct at two ranks higher than the CAMI2 gold standard);
v. Either completely incorrect or Kraken2 did not classify the read (index=9, the maximum number of ranks away from the gold standard classification).

Note that the taxonomic assignments given here are purely for illustration purposes, and not indicative of actual taxonomic assignments given. The indices for all reads within a sample were then combined to give the proportion of reads that were correctly classified at the CAMI2 gold standard rank, the proportion of reads that Kraken2 classified at a lower rank than the CAMI2 gold standard, the proportion of reads that Kraken2 classified at a higher rank than the CAMI2 gold standard, the proportion of reads that were classified incorrectly, the proportion of reads that were unclassified, the mean overall index, the mean index for all reads that Kraken2 classified at a higher rank than the CAMI2 gold standard and the mean index for all reads that Kraken2 classified incorrectly. This gives an indication of how close the Kraken2 classifications were on average within different sample types. These calculations were carried out for the Kraken2 classifications with a confidence threshold of 0.00 for all Kraken2 databases, and with confidence thresholds between 0.00 and 1.00 at intervals of 0.05 for the RefSeq Complete V205 database.

### 2.6. Script, dataset and database availability

All scripts used for acquiring samples, database construction, taxonomic classification and subsequent analyses can be found at https://github.com/R-Wright-1/kraken_metaphlan_comparison. All databases, processed datasets, taxonomic classifications output from Kraken2 and MetaPhlAn 3, and files generated during the analysis can be found at https://www.dropbox.com/sh/lvlz2wpsssvsrad/AAC-BkJja8LvlDoNDB4qgnHNa?dl=0. We used MetaPhlAn 3 version 3.0.13 throughout. Kraken2 version 2.0.8-beta was used for all analyses aside from those where the number of minimizers was output; outputting the number of minimizers required Kraken version 2.1.2. This was still in development and not yet available as part of a stable release at the time of use so was installed directly from Github. All analyses were carried out using custom R (version 4.1.2) and Python (version 3.9.7) scripts in R Studio (version 2021.09.0) or Spyder (version 5.1.5), using the additional packages deicode [49], ete3 [52], knitr [53, 54], math, matplotlib [55], multiprocessing [56, 57], numpy, os, pandas [58], pickle, reticulate [59], scikit-bio, and SciPy [60].

## 3. Results

### 3.1. Classification of real metagenome datasets

We initially used several human-associated or environmental datasets for which both MGS and 16S data were available (as used previously by Douglas *et al*. [19]) to compare the taxonomic classifications given by Kraken2 (with a confidence threshold of 0.00 or 1.00 and one of eight different databases) and MetaPhlAn 3 or the curated SILVA v138 database for the 16S samples. The proportion of reads within samples that could be classified varies widely depending on whether MetaPhlAn 3 or Kraken2 was used for taxonomic classification as well as which database or confidence threshold was used with Kraken2 (Figs. 1 and S2). The proportion of reads that are classified (by either Kraken2 or MetaPhlAn 3) is much higher for the human-associated (maximum median proportion of reads classified of ∼91%) than environmental datasets (maximum median proportion of reads classified of ∼59%), and the proportion of reads that can be classified in environmental datasets with either MetaPhlAn 3 or Kraken2 with a confidence threshold of 1.00 is less than 1% (Fig. 1). In general, the proportion of reads that can be classified within samples increases with increasing database size. Although MetaPhlAn 3 with the human-associated datasets appears to be an exception to this (a median of ∼48% of reads are mapped with a database size of only 2.8 GB), it should be noted that MetaPhlAn 3 gives an estimated number of reads for each taxon based on the number of reads that are mapped to the marker gene database used (Fig. 1).

**Figure 1.**
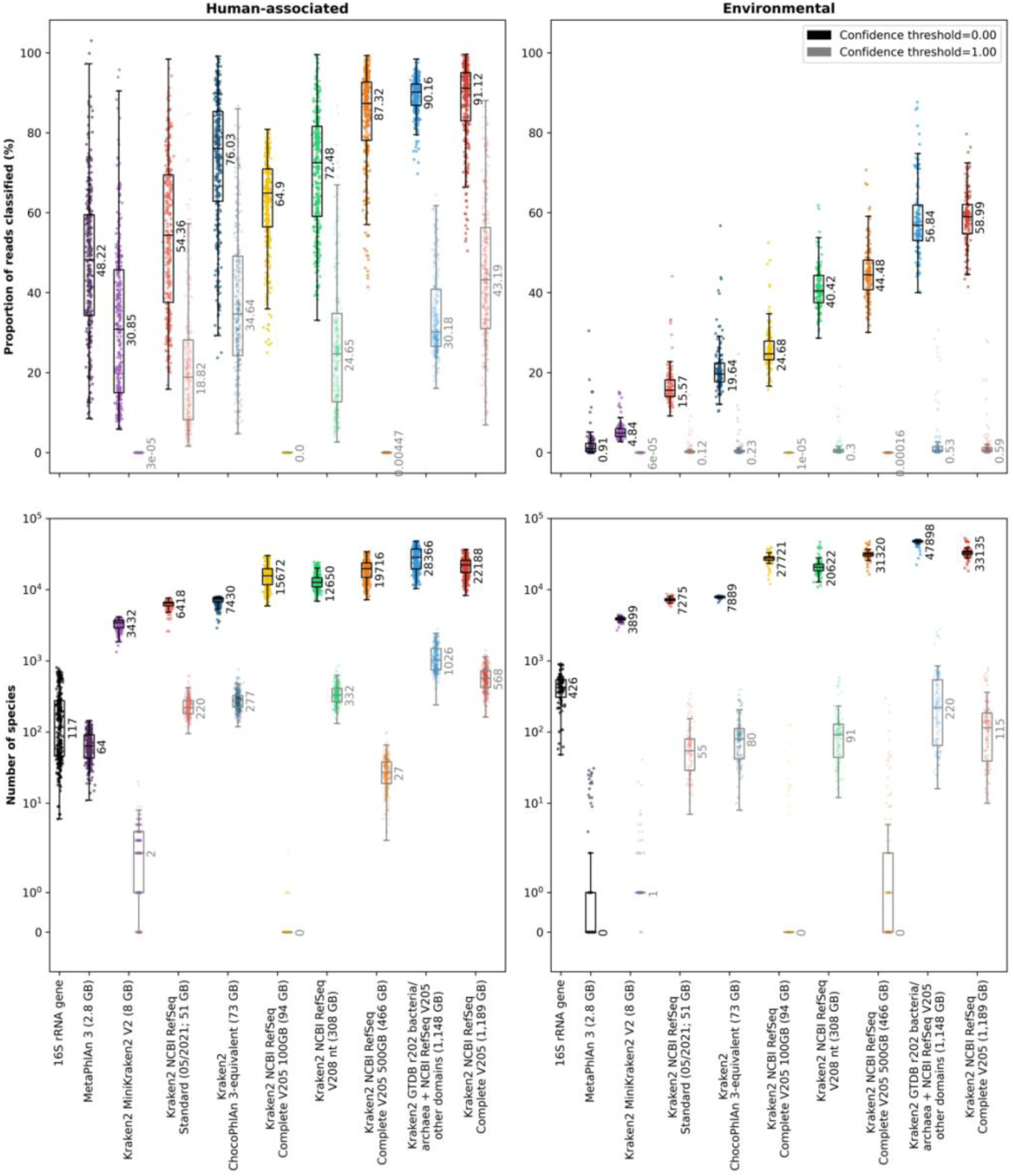
The proportion of reads (MGS samples only; top row) or the number of species classified (16S and MGS samples; bottom row) using different classifiers and databases on real datasets. In each plot, each sample is shown as a single point with boxplots showing the median, upper and lower quartiles while whiskers show the range of the data (1.5 times the interquartile range). Numbers next to boxplots indicate the median for that classifier/database and Kraken2 classifications are shown for confidence thresholds of 0.00 (black boxplots and text and bold colours) or 1.00 (grey boxplots and text and faded colours). Datasets are grouped to either human-associated (left column; Cameroonian, HMP or Indian datasets) or environmental samples (right column; Mammal, Ocean, Primate or Soil [Blueberry] datasets). The disk space used for each of the Kraken2 databases is indicated next to the database name in parentheses. Values for all datasets are shown separately in Fig. S2 alongside the proportion of reads that were classified as bacteria as well as the number of genera that were classified.

As with the proportion of reads classified, the number of species identified within samples also varies widely (across several orders of magnitude) depending on whether MetaPhlAn 3 or Kraken2 was used for taxonomic classification, as well as which database or confidence threshold was used with Kraken2 (Figs. 1 and S2). There were also large variations in the number of species identified between 16S and MGS taxonomic classifications. Both the 16S classifications and Kraken2 classifications with a confidence threshold of 0.00 and all but the smallest databases show that there are, as expected, more species identified within environmental than human-associated datasets (Fig. 1). However, MetaPhlAn 3 and Kraken2 with a confidence threshold of 1.00 both often do not identify any species within environmental samples. Furthermore, even in the human-associated samples, MetaPhlAn 3 only finds a median of 64 species and Kraken2 with the MiniKraken2 V2, NCBI RefSeq Complete V205 100 GB or NCBI RefSeq Complete V205 500 GB databases and a confidence threshold of 1.00 find a median of only 2, 0 or 27 species, respectively. While these all likely represent large underestimations of the true number of species present, Kraken2 with a confidence threshold of 0.00 conversely likely overestimates the number of species present in many cases.

Given the large differences in the proportion of reads classified as well as the number of species identified within human-associated and environmental datasets by both Kraken2 and MetaPhlAn 3, we next turned to simulated and mock samples to determine which tool-parameter-database combination could give us taxonomic classifications closest to those that were observed.

### 3.2. Database characteristics and coverage of the taxa in samples

We first examined whether the databases used covered all species in the mock and simulated samples. We calculated this individually for each sample both in terms of the proportion of species covered and the proportion of the reads in the sample that this equated to. The general trend is for the proportion of species and reads covered by the databases to get larger as database size increases, with the NCBI RefSeq V208 nt and the GTDB r202 bacteria/archaea + NCBI RefSeq V205 other domains database being exceptions in terms of both median and mean, and the ChocoPhlAn 3 and NCBI RefSeq Standard (05/2021) being an exception in terms of the mean (Fig. 2). The NCBI RefSeq Complete V205 database covers the largest proportion of both taxa and reads in each sample, with the median coverage being 99.1% of taxa (mean 98.9%, range 94.3-100%) and 99.9% of reads (mean 99.4%, range 87.2-100%).

**Figure 2.**
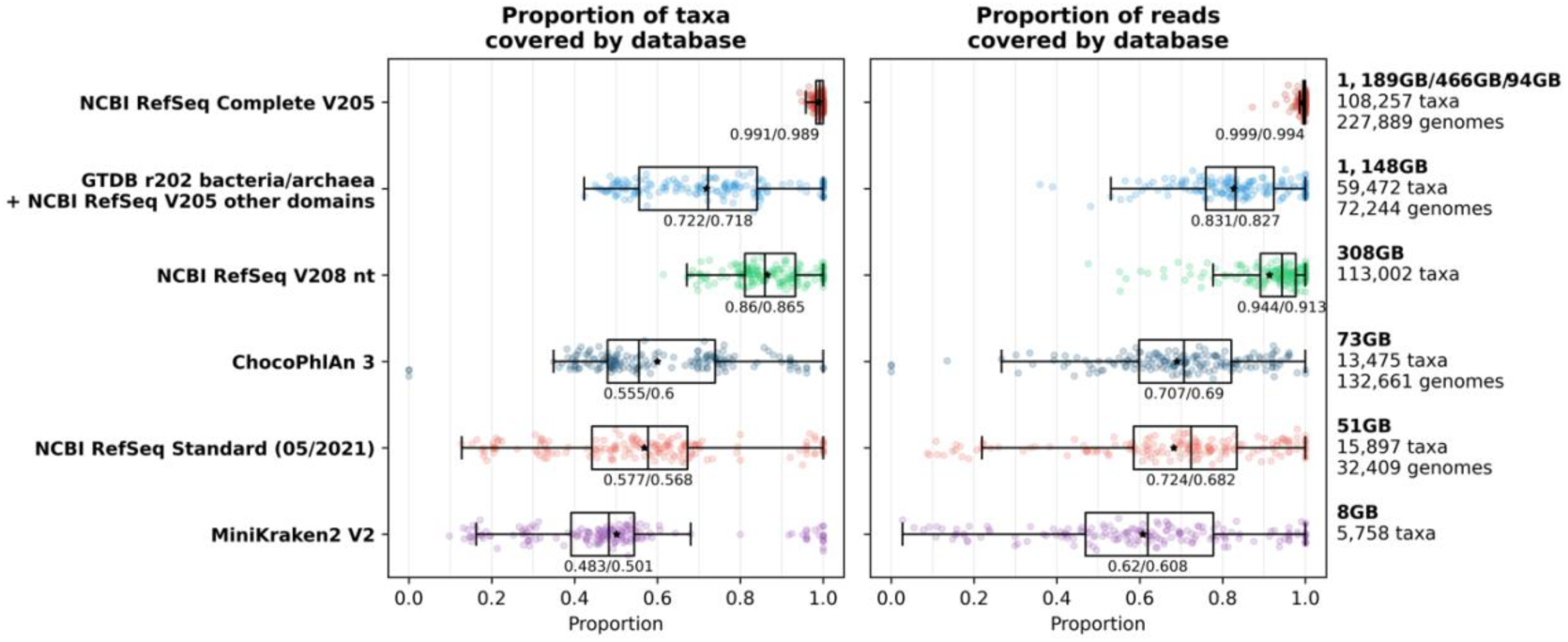
The proportion of taxa (left) and reads (right) within mock and simulated samples that are covered by each database. Each sample is shown as a single point with boxplots showing the median, upper and lower quartiles while whiskers show the range of the data (1.5 times the interquartile range). Black stars indicate the mean for samples and numbers below the boxes show medians/means for all samples within each database. Full descriptions of the taxa contained within each database are given in the Methods section while a summary of the size, number of taxa and number of genomes (where this information is available) contained within each database is given on the right.

### 3.3. Effect of varying Kraken2 confidence threshold on classification accuracy

To determine an appropriate confidence threshold to use, we classified all simulated community samples using Kraken2 followed by Bracken (species level) at each confidence threshold between 0.00 and 1.00 at intervals of 0.05. We then calculated a range of basic (*i*.*e*., precision, recall and F1 score) as well as alpha- and beta-diversity metrics on the compositions as classified by Kraken2 compared with the known composition of the samples. These metrics were calculated for Kraken2 run with all databases using each confidence threshold, but due to the much greater proportion of taxa and reads covered by the NCBI RefSeq Complete V205 database (Fig. 2), we have focused on this database in this section.

#### 3.3.1. Precision, recall and F1 score

We first calculated the precision, recall and F1 score based on either the number of taxa or the number of the reads that were correctly or incorrectly classified, as well as the mean of the taxa and reads for each metric (Figs. 3 and S3). The precision based on the number of reads varied more strongly than that based on the number of taxa, while the opposite was true for the recall (Fig. S3). This meant that the mean precision more closely followed the pattern of the precision based on the number of reads and was maximised at a high confidence threshold (maximum of 0.726 at a confidence threshold of 0.80), while the mean recall didn’t vary as strongly and was maximised at a low confidence threshold (maximum of 0.69 at a confidence threshold of 0.10; Fig. 3). The mean F1 score therefore increased up until a confidence threshold of 0.60 (maximum 0.684) before decreasing (Fig. 3).

**Figure 3.**
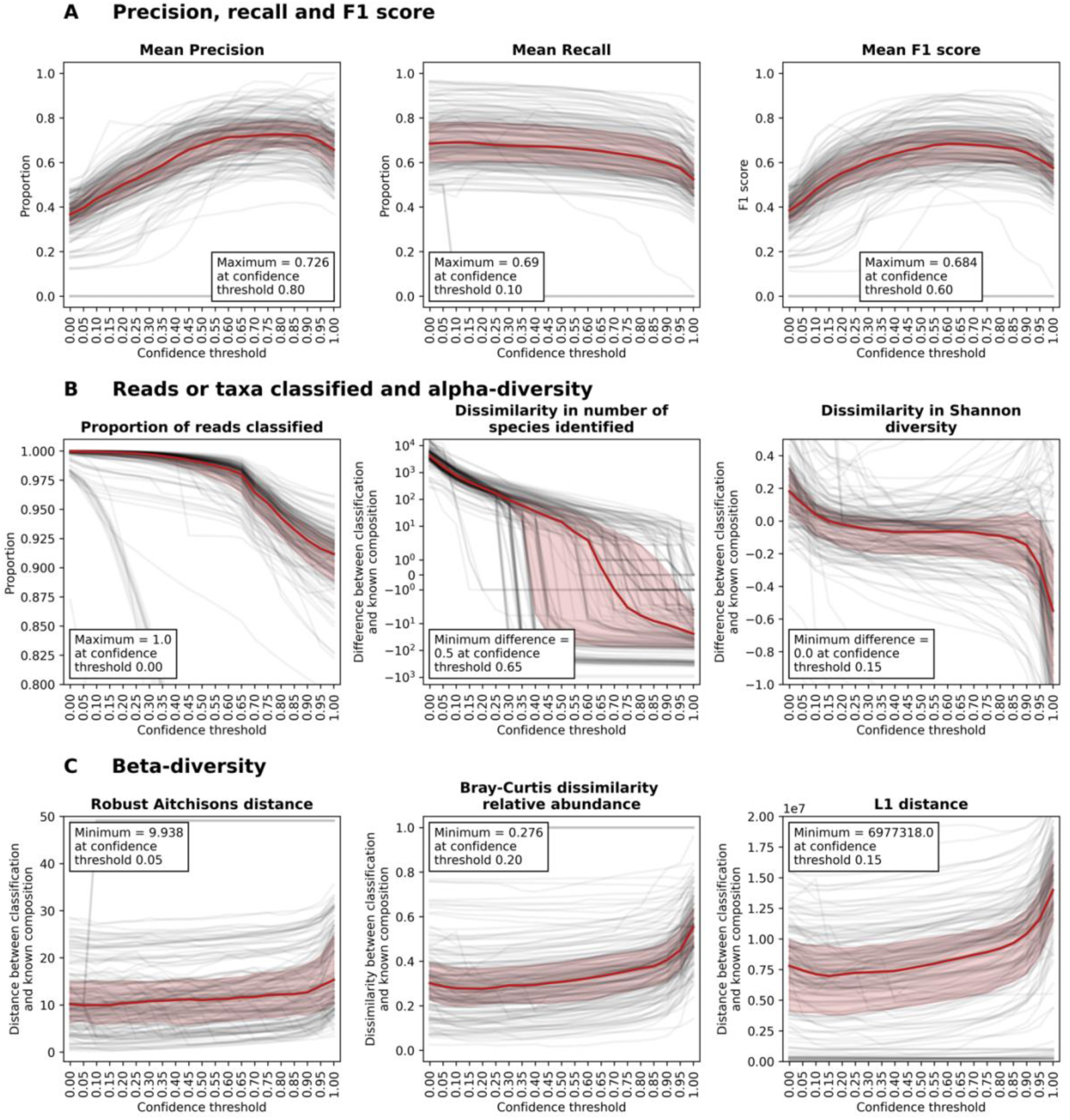
Performance metrics for samples run with Kraken2 against the full database built with NCBI RefSeq Complete V205 with confidence thresholds of 0.00 to 1.00 at intervals of 0.05. In each plot, black lines are for individual samples across different confidence thresholds and red lines and shaded areas show the median and upper and lower quartiles, respectively. Boxes with text indicate the optimal value and confidence threshold for each metric. (**A**) shows precision, recall and F1 score, which are each the mean of the precision, recall and F1 scores based on both the number of taxa and the number of reads classified by Kraken2 as compared with the known sample compositions. These are shown in addition to the means in Fig. S3. (**B**) shows the proportion of reads classified and the dissimilarity in the number of species identified and Shannon diversity between Kraken2-classified samples and the known composition. Positive values show that the metric was higher in Kraken2 classifications than in the known composition, while negative values show the opposite. Additional alpha-diversity metrics are shown in Fig. S4. (**C**) shows the distance or dissimilarity between Kraken2 classifications and the known composition for the Robust Aitchison’s distance, Bray-Curtis dissimilarity calculated with relative abundance values and L1 distance. Additional beta-diversity metrics are shown in Fig. S5.

#### 3.3.2. Proportion of reads classified, number of species identified and alpha-diversity and evenness

We next calculated the proportion of reads classified, the number of species identified and a range of alpha-diversity metrics (Simpson’s, Shannon or Faith’s taxonomic diversity, Chao1 richness or McIntosh’s, Pielou’s or Simpson’s evenness) for both the Kraken2-classified compositions and the known compositions (Figs. 3 and S4). For all of these metrics, aside from the proportion of reads classified, we subtracted the values calculated on the known composition from the values calculated on the Kraken2-classified composition for each sample, meaning that a value of above zero shows that the metric of the Kraken2-classified composition was higher than for the known composition, and a value of below zero shows the opposite. The proportion of reads that were classified decreased with increasing confidence threshold; while some individual samples have different trajectories, the decrease in the median proportion of reads classified is not noteworthy until the confidence threshold is above 0.65 (Fig. 3). The dissimilarity in the number of species classified was initially very high and decreased sharply with increasing confidence threshold, but this decrease started to level off at a confidence threshold of 0.40-0.50 and the median number of species classified was the most similar to the known composition at a confidence threshold of 0.65. All alpha-diversity metrics started off much higher for the Kraken2-classifications than the known composition and decreased with increasing confidence threshold, but the magnitude of these decreases and the confidence thresholds at which they occurred differed (Fig. S4). The Shannon diversity of the Kraken2-classified samples decreased steeply between confidence thresholds of 0.00 and approximately 0.20, the diversity was similar to the known composition between here and confidence thresholds of approximately 0.90 and then again decreased more steeply between here and 1.00. All evenness metrics were lower for the Kraken2-classifications than the known composition at low confidence thresholds. The evenness of the Kraken2-classified samples increased as confidence threshold increases and then at very high confidence thresholds (above 0.90) it either decreased again (Pielou’s or Simpson’s evenness) or sharply increased (McIntosh’s evenness).

#### 3.3.3. Beta-diversity

Finally, we calculated the distance between the Kraken2-classifications and the known compositions using a range of beta-diversity metrics; L1 distance, robust Aitchison’s distance, Aitchison’s distance, Bray-Curtis dissimilarity or weighted and unweighted taxonomic UniFrac distance (Figs. 3 and S5). Only the Aitchison’s and unweighted taxonomic UniFrac distance decreased with increasing confidence threshold, so the minimum distance between the Kraken2-classified and the known compositions were at a confidence threshold of 1.00. All other distances decreased slightly initially with increasing confidence threshold, were minimized at confidence thresholds of 0.05-0.20, and then they increased with increasing confidence threshold.

#### 3.3.4. Choosing an optimal confidence threshold and the metric to base this on

The optimal confidence threshold varies depending on which metric we look at (Figs. 3 and S3-S5) and the metric that is most important for making this choice may vary depending on the research question at hand. When choosing the optimal confidence threshold to use we have mainly focused on the mean F1 score and L1 distance because typically in microbiome research, we are concerned with whether certain taxa are present or absent and whether these differ in abundance, and these metrics mainly capture the dynamics of changing confidence thresholds. We are also interested in whether the taxa that were identified in our samples are closely related to those that are known to be present, and how much the abundances of these differ, *i*.*e*., weighted UniFrac distance. We were, however, unable to find a phylogenetic tree that encompassed all NCBI RefSeq taxa and the computational burden of making such a tree would be too large to carry out here (>100,000 taxa were classified across all samples). Therefore, we calculated our UniFrac distances using taxonomic rather than phylogenetic distance. The median distance using weighted taxonomic UniFrac distance didn’t vary much with increasing confidence threshold until very high confidence thresholds (Fig. S5), and it therefore wasn’t very useful for determining optimal confidence thresholds. In most of our subsequent comparisons we focus on the proportion of reads that are classified, mean F1 score, number of species identified, Shannon diversity and L1 distance. We use the proportion of reads classified, number of species identified and the Shannon diversity to ensure no large deviations from what is expected, while we have used either the maximum mean F1 score or minimum L1 distance to determine the optimal method to use. The optimal confidence threshold for Kraken2 with the NCBI RefSeq Complete V205 database on these simulated samples is therefore either 0.60 (maximum mean F1 score, as well as close to where the number of species classified was the same as in the known composition, minimum Chao1 richness and Simpson’s evenness differences) or 0.15 (minimum L1 distance, as well as minimum weighted UniFrac taxonomic distance, maximum recall and precision based on the reads classified, and minimum Shannon diversity difference).

### 3.4 Effect of different sample characteristics on optimal confidence threshold and classification accuracy

In the Parks *et al*. [8] study, samples were simulated with: (i) varying average nucleotide identity (ANI) to reference genomes (95, 97, 99 or 100%); (ii) low (100) or high (500) average numbers of species per sample; and (iii) with low or high strain diversity within species, with one or up to 10 strains per species, respectively. Parks *et al*. constructed samples with all combinations of these characteristics aside from the 95% ANI and high species diversity (*n*=14 combinations). We used these samples to determine whether these sample characteristics had an impact on the optimal confidence threshold as well as on classification accuracy metrics when using the overall optimal confidence threshold for maximising mean F1 score (0.60).

The different sample characteristics do not have a large impact on what the optimal confidence threshold is for each metric. For example, the optimal confidence threshold for maximising the mean F1 score overall is 0.60 and this only varies between 0.55 and 0.75 for samples with different characteristics (Fig. S6). The highest optimal confidence threshold (0.75) is for samples with an ANI of 100%, low species diversity and no strain diversity, while the lowest optimal confidence threshold (0.55) is found for a few different sample types although generally, for the samples tested here, if species diversity is high and strain diversity is present then the optimal confidence threshold is 0.55. Next, we examined the effect of sample characteristics on classification accuracy at the overall optimal confidence threshold for maximising mean F1 score (0.60; Fig. S7). We find that both the proportion of reads classified and the mean F1 score are both lowest when ANI is 100%, particularly in the absence of strain diversity, while both are highest when ANI is 95% in the absence of strain diversity. This pattern is mainly mirrored with the L1 distance, while the median Shannon diversity tends to be closest to the known composition of samples when species and strain diversity are absent, and furthest from the known composition when both species and strain diversity are present.

### 3.5 Effect of Kraken2 minimizer filtering or filtering taxa based on confidence thresholds on classification accuracy

We investigated filtering taxa based on either the number of distinct minimizers that a taxonomic classification was based on (an experimental feature adapted from KrakenUniq [38]; Figs S8 and S9) or whether they were present at higher confidence thresholds (Fig. S10; see Supplementary Results Section). However, we found that filtering based on the number of minimizers was less effective at increasing precision than altering the confidence threshold was because filtering on the number of minimizers led to a larger decrease in the proportion of reads classified (Fig. S8), likely because the maximum number of distinct minimizers for false and true positive taxa was similar. We also felt that it was more logical to apply the confidence threshold on a per-read basis than filtering based on whether the taxa are identified at all at a particular confidence threshold, although these results are in Fig. S9.

### 3.6 Effect of Kraken2 database choice on classification accuracy

To explore the effect of database choice on classification accuracy we classified all simulated and mock community samples using Kraken2 followed by Bracken for each of eight databases (see Methods for further details on these databases) at all confidence thresholds between 0.00 and 1.00 at intervals of 0.05. We then calculated all metrics on the compositions as classified by Kraken2 and the different databases compared with the known composition of the samples (as detailed above). We carried out these comparisons in order to determine: (i) the impact of confidence threshold on classification accuracy with different databases; (ii) the impact of including more or less taxa in a database, *i*.*e*., the difference between classifications obtained with NCBI RefSeq Complete V205, GTDB r202 bacteria/archaea + NCBI RefSeq V205 other domains, the ChocoPhlAn 3-equivalent and the Standard (05/2021) database; (iii) the impact of including redundancy, *i*.*e*., the difference between classifications obtained with NCBI RefSeq Complete V205 and NCBI RefSeq V208 nt; and (iv) the impact of limiting database size by down-sampling minimizers during the database build, *i*.*e*., the difference between the full, 500 GB and 100 GB size-limited versions of the NCBI RefSeq Complete V205 and the MiniKraken V2 database. Note that the MiniKraken V2 database was built using an older NCBI RefSeq version and includes fewer domains than the databases that we have built, but we included it for comparability with what many researchers may use given its easy accessibility and small size.

#### 3.6.1. Impact of confidence threshold on classification accuracy with different databases

As confidence threshold is increased, we see two different patterns in how each metric changes depending on whether the database has been limited by down-sampling of minimizers or not (Fig. 4). If minimizers have been down-sampled then: the proportion of reads that are classified decreases rapidly; the mean F1 score increases slightly initially and then decreases, with a small second peak; the Shannon diversity sharply decreases initially and then stays relatively consistent until almost no reads are classified; and the L1 distance decreases slightly initially and then increases until almost no reads are classified. The other databases that have not been down-sampled show similar patterns to that described above for the NCBI RefSeq Complete V205. Regardless of whether minimizers have been down-sampled or not, the number of species identified decreases sharply initially with increasing confidence threshold. The number of species identified quickly drops below the number of species in the known composition for the databases where minimizers have been down-sampled. Interestingly, for the databases that have not been down-sampled, the NCBI RefSeq Complete V205 is the only database where the number of species identified is ever below the number of species in the known composition, even though this database has more taxa than any other database (Fig. 2), presumably because a given taxon is more likely to be present in the database, reducing erroneous classifications to other similar organisms. The optimal confidence threshold to use based on either mean F1 score or L1 distance varies depending on the database used: for the down-sampled databases it is 0.05-0.30 based on mean F1 score or 0.00-0.10 based on L1 distance, and for the other databases it is 0.45-0.90 based on mean F1 score and 0.10-0.40 based on L1 distance.

**Figure 4.**
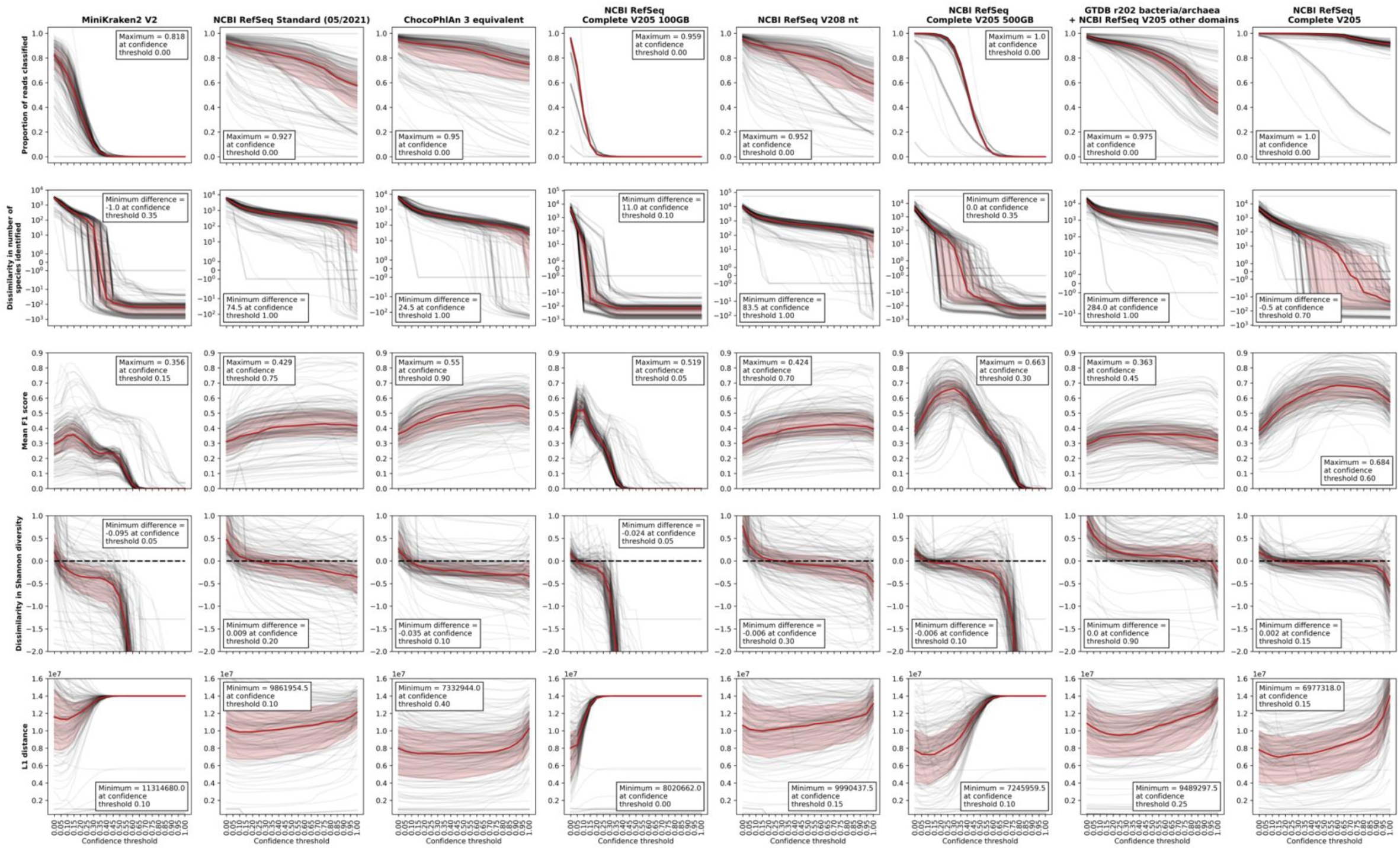
Performance metrics for samples run using Kraken2 with each of eight different databases (columns). Databases are sorted by increasing size (left to right; see Fig. 2). The proportion of reads classified, dissimilarity in the number of species identified, mean F1 score, dissimilarity in Shannon diversity and L1 distance (rows) are shown for all samples across confidence thresholds between the Kraken2-classified samples and the known composition. In each plot, black lines are for individual samples across different confidence thresholds while red lines and shaded areas showing the median and upper and lower quartiles, respectively. Boxes in plots show the value of that metric when it is optimised (maximised for proportion of reads classified and mean F1 score or minimised for the number of species identified, Shannon diversity and L1 distance) and the confidence threshold at which this occurs.

#### 3.6.2. Impact of the number of taxa included in the database on classification accuracy

When we compare the databases that have not been down-sampled, but that have different numbers of taxa (Fig. 4), the NCBI RefSeq Complete V205 has the largest maximum number of reads classified and mean F1 score as well as the smallest minimum L1 distance, while the GTDB r202 bacteria/archaea + NCBI RefSeq V205 other domains has the smallest Shannon diversity difference (although all are reasonably similar for Shannon diversity). Although the NCBI RefSeq Complete V205 database is the largest database of all tested and has the highest mean F1 score (0.684), the maximum mean F1 score for the other databases does not follow the same logic; the ChocoPhlAn 3-equivalent database (73 GB) had a higher maximum mean F1 score (0.55) than the Standard database (0.429; 51 GB), while the GTDB r202 bacteria/archaea + NCBI RefSeq V205 other domains (1,148 GB) had the lowest mean F1 score (0.363). A similar pattern was observed for the minimum L1 distance, with the NCBI RefSeq Complete V205 database having the lowest, followed by the ChocoPhlAn 3-equivalent and then the GTDB r202 bacteria/archaea + NCBI RefSeq V205 other domains and Standard databases.

#### 3.6.3. Impact of including redundancy in a database on classification accuracy

When we compare the NCBI RefSeq Complete V205 with the NCBI RefSeq V208 nt (non-redundant) database, the NCBI RefSeq Complete V205 out-performs the non-redundant database in almost every instance (Fig. 4). The optimised values for the NCBI RefSeq Complete V205 are always closer to the known composition than the non-redundant database. However, there is less variation with increasing confidence threshold for the non-redundant database and, at the highest confidence thresholds, the L1 distance between the NCBI RefSeq Complete V205-classified samples and the known composition is higher than for the non-redundant database.

#### 3.6.4. Impact of database size limitation by minimizer down-sampling on classification accuracy

When examining the size-limited databases with the NCBI RefSeq Complete V205 database, we see that they always perform worse than the complete database and this decrease in performance is in line with the extent to which down-sampling has been performed (Fig. 4). However, the NCBI RefSeq Complete V205 500 GB database is the second-best performing database of all other databases tested in terms of both optimal mean F1 score and L1 distance, despite being approximately 2.5 times smaller than the NCBI RefSeq Complete V205 database.

#### 3.6.5. Choice of database to use with Kraken2

The NCBI RefSeq Complete V205 database always out-performs the other databases that we have tested with Kraken2, however, the computational requirements required to use this database may make this infeasible for many researchers. This section shows that if a researcher is unable to use the largest database, then the next best options are either to use the NCBI RefSeq Complete V205 500 GB or to use a smaller, curated database, such as the one that we have made with the taxa that are included in ChocoPhlAn 3. Excessive size limitation leads to extremely poor performance that cannot be improved by increasing the confidence threshold and the user must treat the use of confidence thresholds differently depending on whether the database is size-limited with down-sampled minimizers or not. If using a database with down-sampled minimizers, then the confidence threshold can be increased to between 0.05 or 0.30 (depending on the extent to which the minimizers have been down-sampled), but above this too few reads are classified, and performance suffers. If using a database that is not size-limited, then the confidence threshold can be increased without the number of reads classified decreasing too drastically. If using a smaller database, like the ChocoPhlAn 3-equivalent database, then the confidence threshold can be increased even further than with the NCBI RefSeq Complete V205 database. For a modest size increase between the Standard (51 GB) and ChocoPhlAn 3-equivalent (73 GB) database, the ChocoPhlAn 3-equivalent database outperforms the Standard database in terms of both mean F1 score and L1 distance. We do not recommend using the MiniKraken2 V2 database; using this database leads to substantially worse classifications than with any other database.

### 3.7. Kraken2 *vs* MetaPhlAn 3

In this section we compare the optimal MetaPhlAn 3 parameters with the optimal confidence thresholds for Kraken2 run with either the ChocoPhlAn 3-equivalent database or the NCBI RefSeq Complete V205 database. We performed the above comparisons using different parameters and databases for Kraken2 because there are limited recommendations given by the tool creators, but MetaPhlAn 3 has recommended settings and default parameters. While we did investigate different MetaPhlAn 3 parameters (Fig. S11 and Supplementary Results Section), these did not lead to large differences in performance, as they did for Kraken2. We therefore chose to compare the default MetaPhlAn 3 --very-sensitive Bowtie 2 option, as well as with the slightly less stringent --sensitive option, both with estimation of the number of reads contributed by each taxon (using the default statistical method), with Kraken2 run with the NCBI RefSeq Complete V205 and the ChocoPhlAn 3-equivalent databases, as well as several confidence thresholds (0.00, 0.50 and 1.00 as well as those for the optimal mean F1 score or L1 distance for each database).

#### 3.7.1. Comparison of key performance metrics at the read level

Kraken2 (regardless of whether it is run with the NCBI RefSeq Complete V205 or ChocoPhlAn 3 database) classifies a higher proportion of reads than MetaPhlAn 3 (Fig. 5). These classifications are also carried out on a read-by-read basis, rather than a prediction based on identifying a set of core marker genes, so this proportion is based on the actual number of reads classified rather than a predicted portion of reads. The median number of species identified is always too low for MetaPhlAn 3 (difference of -71.5) and too high for Kraken2 with the ChocoPhlAn 3 database (difference of 28 to 6,096), while Kraken2 with the NCBI RefSeq Complete V205 database (difference of -22 to 4,250) is closest to the real median number of species at a confidence threshold of 0.60 (of those shown here; difference of 4). The mean F1 scores for Kraken2 run with the NCBI RefSeq Complete V205 database at confidence thresholds of 0.50, 0.60 or 1.00 are higher than for MetaPhlAn 3, however, the mean F1 score for a confidence threshold of 0.15 is in between the mean F1 scores for MetaPhlAn 3 estimated reads with the very-sensitive or sensitive options and is lower than either MetaPhlAn 3 option for a confidence threshold of 0.00. These differences are due to the recall (taxa or reads) being much higher for Kraken2 with the NCBI RefSeq Complete V205 than for either MetaPhlAn 3 option, while the precision (based on taxa) for Kraken2 with the NCBI RefSeq Complete V205 is very low at low confidence thresholds and is only higher than MetaPhlAn 3 when the confidence threshold is 1.00 (Fig. S12). For Kraken2 with the ChocoPhlAn 3 database, the highest mean F1 score (0.55), which is at a confidence threshold of 0.90, is marginally higher than the mean F1 scores for MetaPhlAn 3 estimated reads with either the sensitive (0.54) or the very-sensitive Bowtie2 settings (0.51). Recall (taxa or reads) for Kraken2 with the ChocoPhlAn 3 database is always higher while the precision (taxa or reads) is always lower than for MetaPhlAn 3 with either Bowtie2 setting. The Shannon diversity for classified samples is closer to the true diversity for all Kraken2 classifications, aside from with the NCBI RefSeq Complete V205 database at a confidence threshold of 1.00, than it is for MetaPhlAn 3. Likewise, in almost all cases, the L1 distances obtained using Kraken2 are lower than for MetaPhlAn 3; the only exceptions to this are when the confidence threshold is set to 1.00.

**Figure 5.**
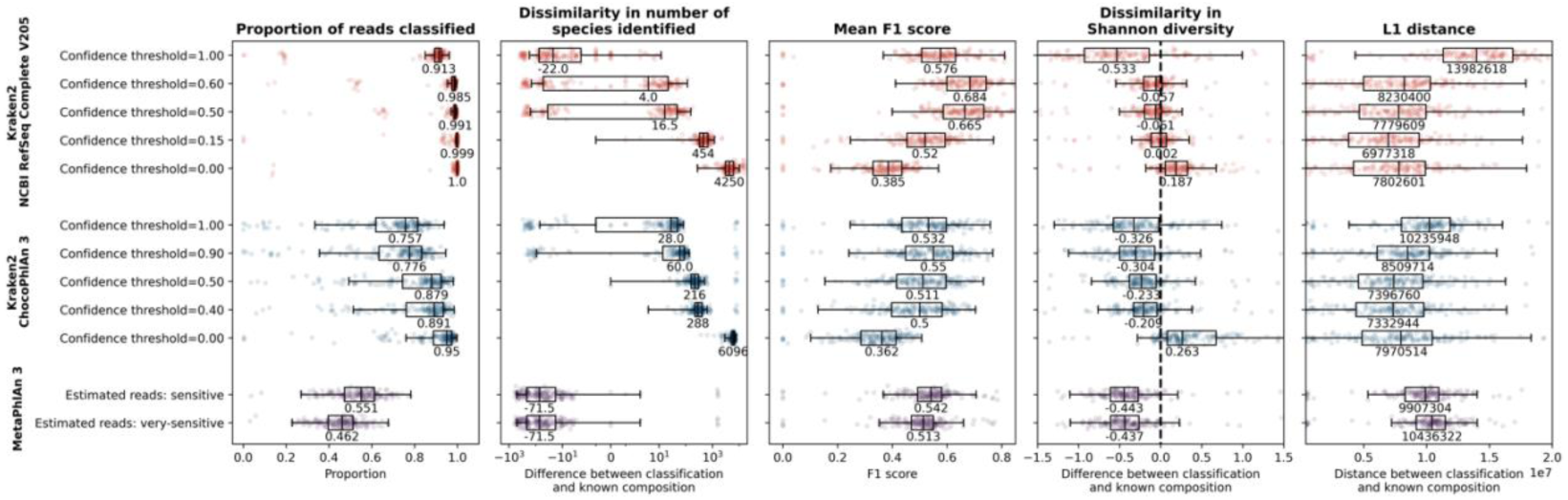
A comparison of the performance of MetaPhlAn 3 and Kraken2 with either the ChocoPhlAn 3 or the NCBI RefSeq Complete V205 databases, showing the proportion of reads that are classified (left), the dissimilarity in the number of species identified (second from left), the mean F1 score (middle), the dissimilarity between the classified and the known Shannon diversity (second from right) and the L1 distance between the classified and the known compositions (right). In each plot, each sample is shown as a single point with boxplots showing the median, upper and lower quartiles while whiskers show the range of the data (1.5 times the interquartile range). Kraken2 run with the NCBI RefSeq Complete V205 databases (top) is shown in red, Kraken2 run with the ChocoPhlAn 3 database (middle) is shown in blue and MetaPhlAn 3 (bottom) is shown in purple. For the two Kraken2 databases we have shown the results for running with confidence thresholds of 0.00, 0.50 and 1.00 in addition to the threshold at which the mean F1 score (0.60 for NCBI RefSeq Complete V205 and 0.90 for ChocoPhlAn 3) or L1 distance (0.15 for NCBI RefSeq Complete V205 and 0.40 for L1 distance) was optimised. For MetaPhlAn 3 we have shown the default estimated reads (using the very-sensitive Bowtie2 option) as well as the option where the L1 distance was optimised and the mean F1 score was almost optimised (the estimated reads with the sensitive Bowtie2 option).

#### 3.7.2. Accuracy at the community level

The analyses that we have shown thus far have all been based on a known composition that includes the number of reads contributed by each taxon. Next, we investigated the 11 samples for which the known composition was provided as either the proportion of taxa in the original community, or as the proportion of DNA from each organism that was used for sequencing. We do note, however, that there are limitations to this, including: (1) these samples only have a maximum of 22 species (median of nine), and are therefore not representative of most environmental samples; (2) all but two of the samples were made up of an even mixture of all taxa, and the two that were not were both ZymoBIOMICS Microbial community standards (from different batches and prepared/sequenced by different facilities); (3) we were not able to find the genome size for all taxonomy IDs in our classified samples: 2,793 of 25,126 taxonomy IDs did not have a genome size, although this accounted for only a median of 0.004% of reads (maximum 0.54%); (4) we were not able to find details of genome copy number (or average genome copy number) for all organisms/genomes in our database; and (5) the small sample numbers (*n*=11) compared with the above comparisons on the number of reads (*n*=164).

On these samples, MetaPhlAn 3 has a much higher precision (based on the taxa classified) than even Kraken2 with a confidence threshold of 1.00, while Kraken2 (at any confidence threshold) has a higher recall than MetaPhlAn 3 (Fig. S13). MetaPhlAn 3 also has a much higher F1 score than Kraken2, although it is further from the known sample composition in terms of Shannon diversity for confidence thresholds of 0.00, 0.50 and 0.60 and has a higher L1 distance than Kraken2 with a confidence threshold of 0.15. It is noteworthy, however, that the recall, precision and F1 scores based on the taxa here differ a lot from the above comparisons on samples with a known composition at the read level. The MetaPhlAn 3 and Kraken2 recall based on taxa classified was 0.45 and 0.66-0.76 (Fig. S12), respectively, on the above comparisons with known compositions in terms of number of reads and are 0.68 and 0.85-1.0 (Fig. S13), respectively, with known compositions in terms of taxa proportions. This large increase in recall for both MetaPhlAn 3 and Kraken2 is likely due to the reduced number of species within each mock sample here, as we wouldn’t have expected the recall and precision based on the taxa classified to differ between these two comparisons of compositions that were known in terms of number of reads or taxa proportions.

We also investigated the impact of normalising the Kraken2 classifications with genome size, although this had very little impact on the classification accuracy (Fig. S13). Here we divided the number of reads classified by Kraken2 for each taxon by the genome size, prior to conversion to relative abundances. The precision, recall and F1 score (based on the taxa classified) are obviously not expected to vary here, but the differences in Shannon diversity and L1 distance were also minimal. When genome size normalization was carried out, the Shannon diversity and L1 distance were slightly closer to the known composition for confidence thresholds of 0.50 and above than when it was not, while for confidence thresholds below 0.50, not carrying out genome size normalization gave compositions slightly closer to the known composition.

#### 3.7.3. Time and resources required to run

Kraken2 was faster on average at classifying mock or simulated samples when the database was already loaded onto a RAM disk (70 s or 35 s for the NCBI RefSeq Complete V205 or ChocoPhlAn 3 databases, respectively) than MetaPhlAn 3 (4 min 28 s), however, the maximum memory usage for Kraken2 was much greater (956 GB or 72 GB for the NCBI RefSeq Complete V205 or ChocoPhlAn 3 databases, respectively) than MetaPhlAn 3 (2.8 GB; Table S3). Also, the database that we have used here with MetaPhlAn 3 was the default one and – although we did not time these steps – this took only a few minutes to download and install, whereas the databases for Kraken2 took several days to a week to download and build, and this would not be feasible on a server that had less than approximately 1.5 TB of RAM. Using only any available pre-built databases would limit this to only the time needed to download the database, however, this would still be considerably longer for Kraken2 if the NCBI RefSeq Complete V205 database was used. All times and memory usage given here were computed using 24 threads.

### 3.8 Classification of samples simulated to imitate different environments

While many of the simulated or mock samples used above were built using genomes that were already present in databases and have unambiguous taxonomic classifications, the simulated samples created for the Critical Assessment of Metagenome Interpretation challenges (CAMI [5] and CAMI2 [6]) used newly sequenced genomes that do not necessarily have taxonomic classifications at the species level. The provenance of all reads – including the genome that the read came from, the taxonomic classification of the genome and the OTU that the genome was assigned to – as well as a “gold standard” taxonomic profile (described in the methods section) was available for the CAMI2 and both the CAMI and CAMI2 samples, respectively. Using these classifications, the simulated samples had a median of 0-93.6% of reads with species-level taxonomic classifications in the read-by-read taxonomic affiliations and 87.2-100% of reads with species-level taxonomic classifications in the gold standard taxonomic profiles (Fig. S14). Because of the difficulty in assessing differences in precision, alpha- and beta-diversity when not all reads are accounted for in the known composition, we calculate only a few metrics for classifications given by Kraken2 with each of the eight databases, as well as MetaPhlAn 3 with the default estimated reads setting. We then focus on the accuracy of Kraken2 classifications on a read-by-read basis and we developed an index metric to do so.

#### 3.8.1. Proportion taxa and reads covered by databases

We examined the proportion of species in the gold standard sample profiles that were present in the databases as well as the proportion of the reads in the samples that this equated to. As with the other simulated and mock samples (Fig. 2), there was a general trend for the proportion of species covered by the database to get larger as the database size increased when all samples are examined together (Fig. S15). This is mainly due to this being the trend for the mouse gut (*n*=49) and strain madness (*n*=100) datasets, while taxa within the marine (*n*=10), rhizosphere/plant-associated (*n*=21) and original CAMI challenge (*n*=10) samples were not well covered by several of the databases. In these datasets there are circular elements (viruses and plasmids), and the rhizosphere/plant-associated samples additionally have fungal and plant host material present: most of these groups are not present in many of the smaller databases, or are not well represented, explaining why these databases contain a lower proportion of the taxa. Furthermore, some of the NCBI taxonomy IDs that are not present in the ChocoPhlAn 3 or GTDB r202 bacteria/archaea + NCBI RefSeq V205 other domains belong to species with only a single representative genome or a genome with an incomplete assembly, which would not be included in these curated databases. The trend was much the same for the proportion of reads within samples that were covered by databases, with the exception that for the strain madness samples, although the proportion of taxa covered was often low, the proportion of reads covered was not so low. As previously (Fig. 2), the proportion of taxa and reads that were covered by the NCBI RefSeq Complete V205 database was equal to or greater than all other databases.

We also looked at the proportion of taxa in the read-by-read taxonomic profiles at both the species and genus levels that were covered by each database. At the species level, the proportion of taxa covered in most datasets followed similar patterns to those seen in the gold standard profiles: only the GTDB r202 bacteria/archaea + NCBI RefSeq V205 other domains and ChocoPhlAn 3 databases did not perform well on marine samples; only the NCBI RefSeq Complete V205 and NCBI RefSeq V208 nt databases did perform well on the rhizosphere/plant-associated samples; and the performance of the databases generally followed with their size on the strain madness samples, with the GTDB r202 bacteria/archaea + NCBI RefSeq V205 other domains performing particularly poorly (Fig. S16). Because close to zero of the reads in the mouse gut samples had a taxonomic classification at the species level (Fig. S14), these all show that close to 0% of taxa and reads are covered by the databases (Fig. S16). At the genus level, all databases covered identical proportions of both taxa and reads in the strain madness samples and the three largest databases performed identically on the mouse gut samples, with the other databases covering proportions of both taxa and reads in line with their sizes. For the rhizosphere/plant-associated samples, only the NCBI RefSeq Complete V205 and NCBI RefSeq V208 nt datbases covered >90% of taxa in the samples, with the other databases covering 52-68% of taxa, but the proportion of reads covered was >85% for all databases. For the marine samples, the proportion of taxa covered was >90% and the proportion of reads covered was above 95% for all databases aside from the ChocoPhlAn 3-equivalent database.

#### 3.8.2. Proportion of reads classified and recall for different classification tool-database-confidence threshold combinations

The proportion of reads classified as well as recall based on either taxa or reads classified varied strongly with both sample type and classification tool-database combination (Fig. S17). The only classification tool-database combinations that always classified above 95% of the reads in the gold standard profile were Kraken2 with either the NCBI RefSeq Complete V205, the GTDB r202 bacteria/archaea + NCBI RefSeq V205 other domains or NCBI RefSeq Complete V205 100GB databases, while the only sample group where all classification tool-database combinations classified almost all reads was the CAMI2 strain madness group. As for the previous simulated samples (Figs. 4 and 5), Kraken2 with the NCBI RefSeq Complete V205 database generally had the highest recall (based on either taxa or reads classified), with the only notable exception here being for the CAMI2 strain madness samples, where Kraken2 with the NCBI RefSeq Complete V205 database had a recall based on the reads classified of 0.666 while the highest was 0.759 for MetaPhlAn 3 and 0.753 for the best performing Kraken2 database (Standard 05/2021; Fig. S17).

As the confidence threshold was increased, the proportion of reads classified, and recall based on either taxa or reads classified by Kraken2 with each of the eight databases followed similar patterns to those seen above for the other simulated samples (Fig. 4): the databases that were size-limited by down-sampling of minimizers showed steep decreases in all three metrics even at low confidence thresholds (Fig. S18). The decrease in the proportion of reads classified by the databases that were not size-limited was almost linear as the confidence threshold increased, while the recall based on the taxa classified did not change a large amount throughout aside from for the strain madness samples. The recall based on the reads classified was initially relatively consistent but started to decline as certain confidence thresholds were reached, which depended both on the database used as well as the sample type and was generally lower for the strain madness and mouse gut samples than the marine, rhizosphere/plant-associated and first CAMI samples. The recall based on taxa was typically highest for marine samples, with the mouse gut and strain madness samples taking intermediate values, while the recall based on the number of reads was high for the mouse gut and strain madness samples but lower for the marine and rhizosphere/plant-associated samples.

#### 3.8.3. Accuracy on a read-by-read basis

For the CAMI2 samples (marine, rhizosphere/plant-associated, mouse gut and strain madness sample groups), information on the provenance of individual reads was given so we were able to assess the accuracy of Kraken2 classifications on a read-by-read basis. To do this, we developed an index system (described in detail in the methods) whereby an index of 0 indicated that the Kraken2 taxonomic classification was the same as the CAMI2 taxonomic classification (at the lowest rank given by CAMI2), and an index of above 0 indicated the difference between the Kraken2 and CAMI2 taxonomic classifications in number of taxonomic ranks. So, for example, an index of 2 would indicate that the Kraken2 classification was correct at two ranks higher than the CAMI2 gold standard classification. The mean of indices for all reads within a sample is taken to give the overall index for each sample, and this is also calculated separately for only the reads that Kraken2 classified at a higher taxonomic rank than the CAMI2 taxonomic classification and those that Kraken2 classified incorrectly at the rank of the CAMI2 taxonomic classification. This part of the analysis was carried out on the Kraken2 classifications with all eight databases with a confidence threshold of 0.00 and for Kraken2 run with the NCBI RefSeq Complete V205 database using confidence thresholds between 0.00 and 1.00 at intervals of 0.05.

#### 3.8.4. Accuracy on a read-by-read basis for Kraken2 with all databases and no confidence threshold

The NCBI RefSeq Complete V205 database classifications were, on average, closest to those given by CAMI2 (lower median index) than those for any other database in all sample groups aside from the strain madness group (Fig. S19). This is due to a combination of having a relatively low proportion of reads that are either unclassified or incorrectly classified, and a relatively high proportion of reads that are either correctly classified at the rank given by CAMI2 or were only classified to a higher rank than the CAMI2 classification. Interestingly, when reads were classified incorrectly by Kraken2 with the NCBI RefSeq Complete V205 database, the classifications given were further away from the CAMI2 classifications than for the smaller databases (higher median index of reads classified incorrectly). Although the proportion of reads that fell into this category was low, it is likely indicative of conserved/repetitive genomic regions that may match multiple taxa and these having a higher chance of being further from the taxon they should be in a larger database. The NCBI RefSeq Complete V205 100GB database performed particularly poorly on the marine and strain madness datasets while the ChocoPhlAn 3-equivalent database also performed poorly on the marine samples, with both having low proportions of reads correctly classified at the CAMI2 rank and a high overall index on these samples. The ChocoPhlAn 3-equivalent database covers a lower proportion of the taxa in the marine dataset than the other similarly-sized databases (Fig. S16) because several of the genomes in the marine samples are classified as taxa that have only a single genomic representative or the genomes are incomplete, and thus wouldn’t be captured by the clade-specific marker gene approach used to curate this database. For the NCBI RefSeq Complete V205 100GB database, the issues seen for the NCBI RefSeq Complete V205 database are likely exacerbated by excessive down-sampling of minimizers. The GTDB r202 bacteria/archaea + NCBI RefSeq V205 other domains database always classified the highest proportion of reads to a lower taxonomic rank than the CAMI2 classification. It is not possible to definitively determine the reason for this, although we hypothesise that it could be due to the GTDB having a more accurate/complete taxonomic structure for prokaryotes and the lowest common ancestor approach therefore being likely to identify a read in GTDB using the LCA approach to a rank that may not exist in NCBI. In the GTDB, all included prokaryotic genomes have a taxonomic classification at all ranks, whereas in the NCBI RefSeq, genomes may only have, for example, a strain-level and then an order-level taxonomic classification.

For the rhizosphere/plant-associated and the mouse gut samples, the proportion of reads that are not classified and the median index is generally lower for larger databases while the proportion of reads that are correctly classified is higher (Fig. S19). In both sample groups, the MiniKraken V2 database performed the worst, with the indices indicating that classifications were a median of 4.8 or 5.2 taxonomic ranks away from the CAMI2 classifications for the rhizosphere/plant-associated and mouse gut samples, respectively. The marine samples generally had a higher proportion of reads that were correctly classified at the CAMI2 rank and a lower index than other sample types – aside from for the ChocoPhlAn 3-equivalent and NCBI RefSeq Complete V205 100GB databases – likely because a higher proportion of these samples had a species-level classification than for the rhizosphere/plant-associated or mouse gut samples, without having many different strains within each species (Fig. S14). For the strain madness samples, the proportion of reads that were correctly classified tended to get lower and the median index tended to get higher as the Kraken2 database got larger. This is mainly due to the proportion of reads that were classified at a higher rank than that given by CAMI2 getting higher as the database size increases, likely because minimizers within reads are more likely to match multiple strains in the larger and more complete databases. This means that the lowest common ancestor approach used by Kraken2 will likely assign these reads to the species level, or even higher (the median indices for reads classified at a higher rank by Kraken2 ranged between ∼1.3 and ∼3.8 for the GTDB r202 bacteria/archaea + NCBI RefSeq V205 other domains and NCBI RefSeq Complete V205 100 GB databases, respectively).

#### 3.8.5. Accuracy on a read-by-read basis for Kraken2 the NCBI RefSeq Complete V205 database at different confidence thresholds

Finally, we examined how increasing confidence threshold impacted the accuracy of Kraken2 classifications on a read-by-read basis with the NCBI RefSeq Complete V205 database. For all sample types, the proportion of reads that are classified correctly at the rank of the CAMI2 classification, the proportion of reads that are classified incorrectly and the proportion of reads that are classified at a lower rank by Kraken2 than given by the CAMI2 classification decreases as the confidence threshold is increased (Fig. 6). The proportion of reads that are classified at a higher rank by Kraken2 than given by CAMI2 increases initially with increasing confidence threshold and peaks at confidence thresholds between 0.15-0.30 depending on sample type, before decreasing towards zero at a confidence threshold of 1.00. The overall index and the index of reads classified at a higher rank by Kraken2 both increase with increasing confidence threshold for all sample types, while the index of reads that are classified incorrectly decreases initially before increasing with increasing confidence threshold for most sample types, with minima at confidence thresholds of 0.10-0.30 for all sample types except the mouse gut samples. The index of reads classified incorrectly for mouse gut samples only reaches a minimum at a confidence threshold of 1.00.

**Figure 6.**
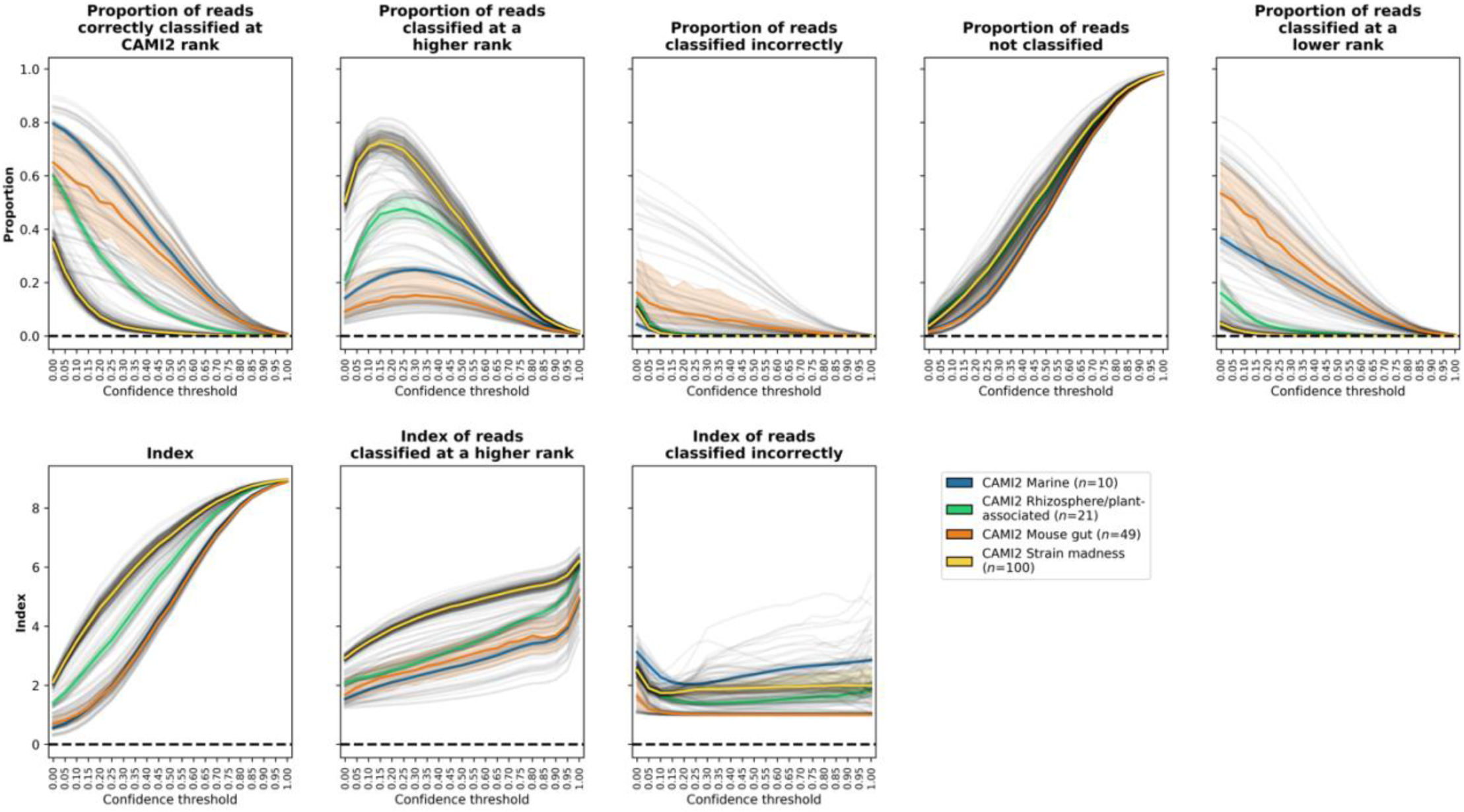
The effect of increasing the confidence threshold with CAMI2 samples classified using Kraken2 with the NCBI RefSeq Complete V205 database on metrics related to the classification of taxa on a read-by-read basis. In each plot, black lines are for individual samples across different confidence thresholds and coloured lines and shaded areas show the median and upper and lower quartiles, respectively, for each sample group.

## 4. Discussion

Here we have used the most recent versions of the two most popular metagenomic taxonomic classification tools, Kraken and MetaPhlAn, to show that using different tools or parameters leads to large differences in both the number of reads classified and species identified within both human-associated or environmental datasets (Fig. 1). To assess which tool-parameter-database combination could give results closest to the ground truth, we tested a range of different database-parameter combinations or parameters with Kraken2 and MetaPhlAn 3, respectively, on a range of simulated and mock samples. We have found that the highest overall performance can be achieved with Kraken2 and a database that contains all genomes in the NCBI RefSeq Complete V205 database, but that setting a confidence threshold within Kraken2 is essential to avoid very low precision (Fig. 5).

There have been several previous studies that include both Kraken (version 1 or 2) and MetaPhlAn (version 2 or 3) as part of a comparison of metagenomic taxonomic classifiers (*e*.*g*. [1–9, 17, 18]). However, these studies have not investigated the use of different confidence thresholds, which we have found to have a large impact on most metrics related to classifier performance (Fig. 2), and very few even set a confidence threshold [1, 3, 17]. These previous studies have also used databases composed of only bacteria [1], only bacterial, archaeal, and viral genomes with complete assemblies [3], a MiniKraken database [1], or do not give full details on what is included in their database [2, 4, 5, 7, 9, 17, 18]. Additionally, while some other studies have used the NCBI non-redundant nucleotide database [6, 8], we are not aware of any studies that have used the full NCBI RefSeq database that we have here, which is likely to have led to significantly worse performance in those previous comparisons (Fig. 4).

Many of these previous studies have also typically not distinguished between methods designed for estimating relative sequence abundance or relative taxon abundance when benchmarking metagenomic taxonomic classifiers [18]. We agree with Sun *et al*. [18] that this may be an important distinction to make and did find that Kraken2 achieved higher performance (more optimal F1 score and L1 distance) on samples with known compositions in terms of the number of reads (sequence abundance; Fig. 5), *i*.*e*., the abundance type for which it was designed. However, this was not necessarily the case for MetaPhlAn 3: on samples with known compositions in terms of the number of cells or DNA extracted (taxon abundance), MetaPhlAn 3 did have a higher F1 score than Kraken2, but it also had a higher L1 distance than for the optimal Kraken2 confidence threshold (Fig. S13). This could be because the samples with known taxon abundance were typically much less complex than those with known sequence abundance, with much fewer taxa and more even compositions, but also, both tools have options for estimating the alternative abundance type, which we have used here (as appropriate). While future metagenomic classifier benchmarking studies should take these differences into account when assessing performance, most researchers conducting microbiome studies should just be aware of which abundance type they are analysing, particularly when comparing their results with other studies.

While we aimed here to systematically investigate the impact of using different databases and parameters on classification accuracy, there are some factors that we didn’t investigate. For example, we did not investigate the impact on classification accuracy of removing eukaryotic genomes or the non-redundant nucleotide sequences (for which the compressed FASTA files and therefore approximate database size reduction totals ∼400 GB) from the NCBI RefSeq Complete V205 database, which could help in reducing the computational burden required for use. While this is something that could be investigated further in the future, other studies have found that only including the organisms of interest in a database can lead to erroneous classifications [10]. We also didn’t investigate the impact of removing low abundance taxa (as has been carried out in other taxonomic classification comparison studies, *e*.*g*. [1]); we felt that applying an appropriate confidence threshold was a better way of dealing with the removal of taxa because, in this way, if a taxon is present but in very low abundances then it should still be identified in the output. It is also worth noting here that there are other tools that previous studies have found to perform better than Kraken2 or MetaPhlAn 3 (*e*.*g*., Microba [8]), however, comprehensively testing all classifier-parameter combinations was outside of the scope of this study.

### Suggestions for use

While the highest performance can be achieved with Kraken2 and the NCBI RefSeq Complete V205 database – and the confidence threshold can be adjusted to optimise the metric of most importance to a researcher – the high computational resource requirement likely makes this infeasible for many researchers. We therefore aim to provide some guidance on how to choose a tool-parameter-database combination to use. If computational resources are limiting and the researcher is not planning to perform other analyses on a read-by-read basis, then the trade-off between size of database, ease of use, memory required, and accuracy of classifications is very good for MetaPhlAn 3 (Figs. 2 and 5 and Table S3). However, if the samples are not of human origin (or possibly just not from Western populations of humans; Fig. S2), it is unlikely that MetaPhlAn 3 will be appropriate (Figs. 1 and S17). In all other circumstances, we recommend using Kraken2; if computational resources are limiting then we suggest choosing the best performing database that resources allow (Fig. 4). In many cases, choosing a confidence threshold to use may mean assessing the trade-off between increasing accuracy vs. the proportion of classified reads that are lost with increasing confidence threshold, *i*.*e*., increasing the confidence threshold as much as possible without leaving too many reads unclassified. The confidence threshold that can be used will therefore depend on characteristics of the sample, such as whether it is of human (increase confidence threshold) or environmental origin (decrease confidence threshold), whether low (increase confidence threshold) or high (decrease confidence threshold) sample diversity is expected, as well as whether the primary concern is minimising false positives (increase confidence threshold) or minimising false negatives (decrease confidence threshold). Performing an initial comparison of different confidence thresholds with a subset of representative study samples may be beneficial for deciding this and the confidence threshold as well as the database used with Kraken2 should be reported within the Methods section of a study.

### Conclusions

We have shown here, using real datasets and simulated or mock samples, that there may not be a one-size-fits-all winner in a comparison between the two most widely used metagenomic taxonomic classifiers, Kraken2 and MetaPhlAn 3. While running Kraken2 with a database containing all genomes in the NCBI RefSeq database achieves the best overall performance, a confidence threshold must be set within Kraken2 to avoid very low precision, and the large amount of memory required to run may make it prohibitive for many researchers. On the other hand, MetaPhlAn 3 does not require many computational resources to run, can be installed and run on samples very quickly and it performs reasonably well out-of-the-box, especially for human-associated samples. The best tool, database or parameter choice for a particular application most likely depends on the scientific question of interest, whether high precision or high recall is more important and what the available computational resources will allow.

## Supporting information

Supplementary figures and results text

## Acknowledgements

The authors gratefully acknowledge funding from an I3V-DMRF Dr. David H. Hubel postdoctoral fellowship for RJW and from an NSERC Discovery Grant 2016-05039 for MGIL. We thank the developers of the MetaPhlAn 3, Kraken2 and Bracken tools as well as the CAMI2 authors for ensuring we were able to access the samples whilst the CAMI2 manuscript was in press and the authors of all studies that made real, simulated or mock community MGS samples publicly available. We also thank Dr. Gavin Douglas for providing processed sequence files for some of the real MGS and 16S samples as well as all members of the Langille Lab for helpful conversations and suggestions throughout the project.

## Author contributions

RJW carried out all bioinformatic and statistical analyses with guidance and technical support from MGIL and AMC. RJW wrote the first draft of the manuscript and all authors contributed substantially to revisions.

## Notes

https://www.dropbox.com/sh/lvlz2wpsssvsrad/AAC-BkJja8LvlDoNDB4qgnHNa?dl=0

https://github.com/R-Wright-1/kraken_metaphlan_comparison

### Competing Interest Statement

The authors have declared no competing interest.

