## Supplementary figures and results text for "From defaults to databases: parameter and database choice dramatically impact the performance of metagenomic taxonomic classification tools"

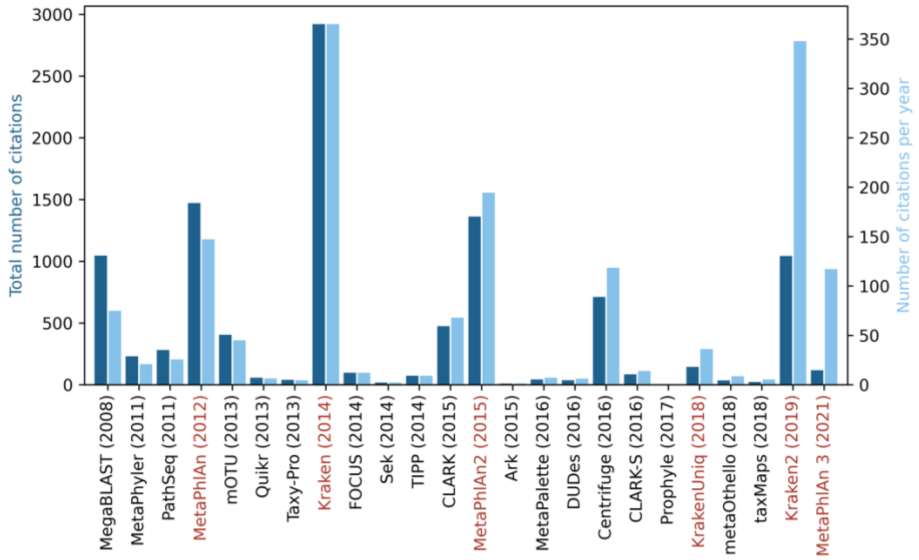


**Figure S1.** Total number of citations (left axis, dark blue) and citations per year (right axis, light blue) for the metagenome taxonomic classifiers included in McIntyre *et al.* [1] and Sczyrba *et al.* [2]. Publications relating to either MetaPhlAn or Kraken are highlighted in red. The numbers of citations for each publication were taken from Google Scholar on March 2$^{nd}$ 2022.

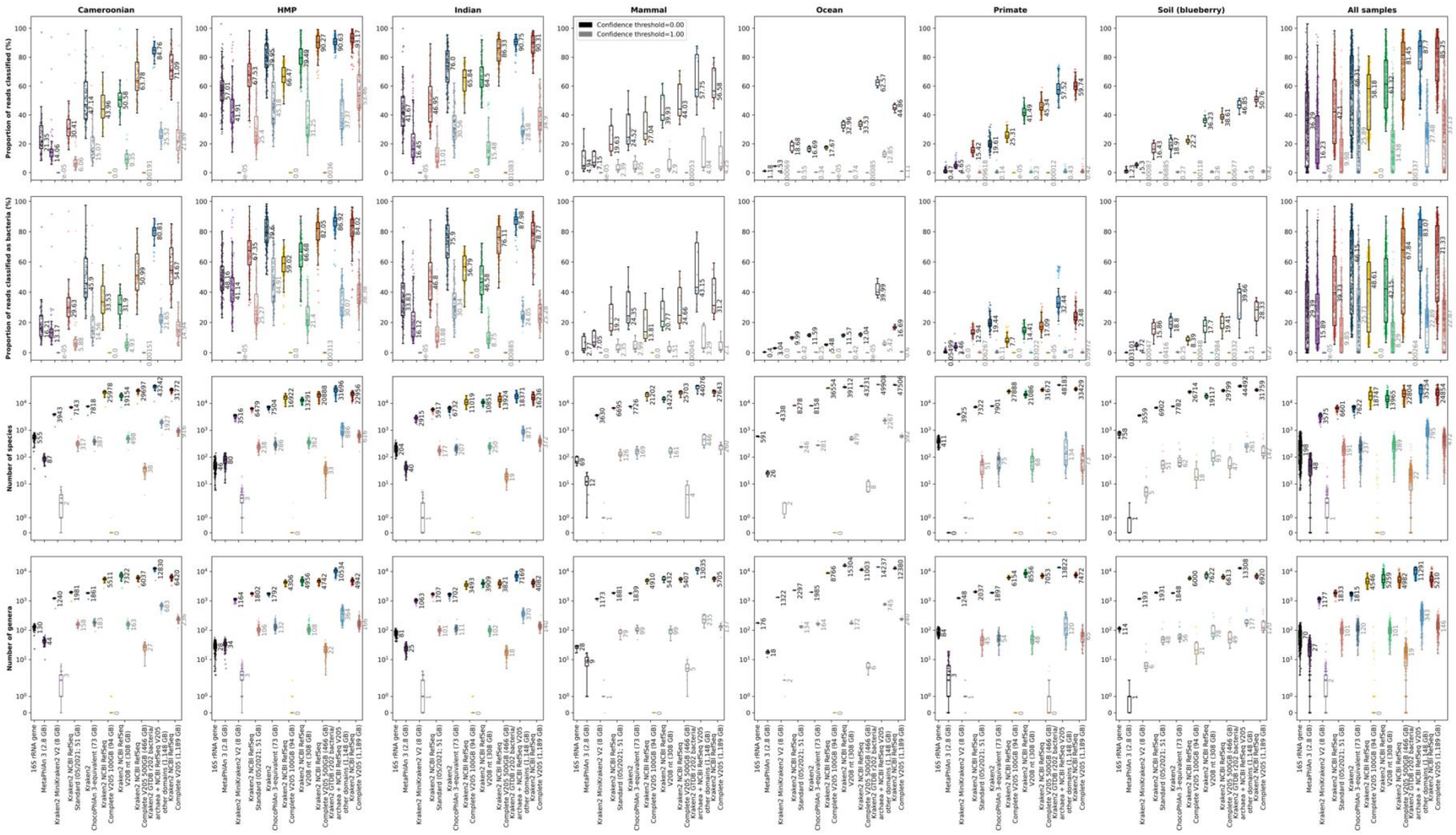


**Figure S2.** The proportion of reads classified (MGS samples only; top row), the proportion of reads classified as bacteria (MGS samples only; second row), the number of species classified (16S and MGS samples; third row) or the number of genera classified (16S and MGS samples; bottom row) using different classifiers and databases on real datasets. In each plot, each sample is shown as a single point with boxplots showing the median, upper and lower quartiles while whiskers show the range of the data (1.5 times the interquartile range). Numbers next to boxplots indicate the median for that classifier/database and Kraken2 classifications are shown for confidence thresholds of 0.00 (black boxplots and text) or 1.00 (grey boxplots and text). Values are shown separately for each dataset (columns) or all together for all samples (final column). The disk space used for each of the Kraken2 databases is indicated next to the database name in parentheses.

8

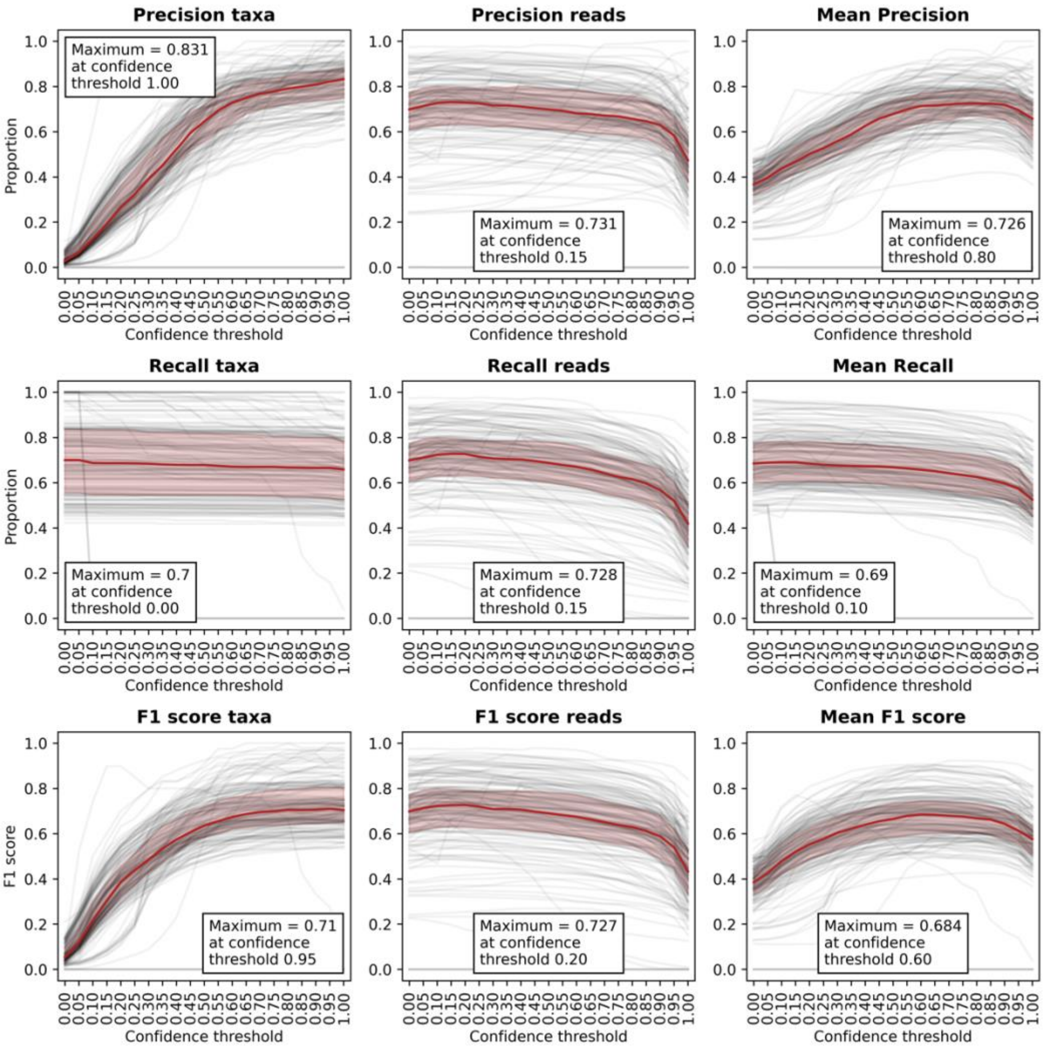


**Figure S3.** Precision, recall and F1 score for samples classified with Kraken2 with the NCBI RefSeq Complete V205 database at confidence thresholds between 0.00 and 1.00 at 0.05 intervals. Here the precision, recall and F1 scores have been calculated for the Kraken2 classifications as compared with the known sample compositions and the means for each of the recall, precision and F1 score are the means of those metrics for the taxa and the reads. In each plot, black lines are for individual samples across different confidence thresholds and red lines and shaded areas show the median and upper and lower quartiles, respectively. Boxes with text indicate the optimal value (maximised metric) and confidence threshold for each metric.

9

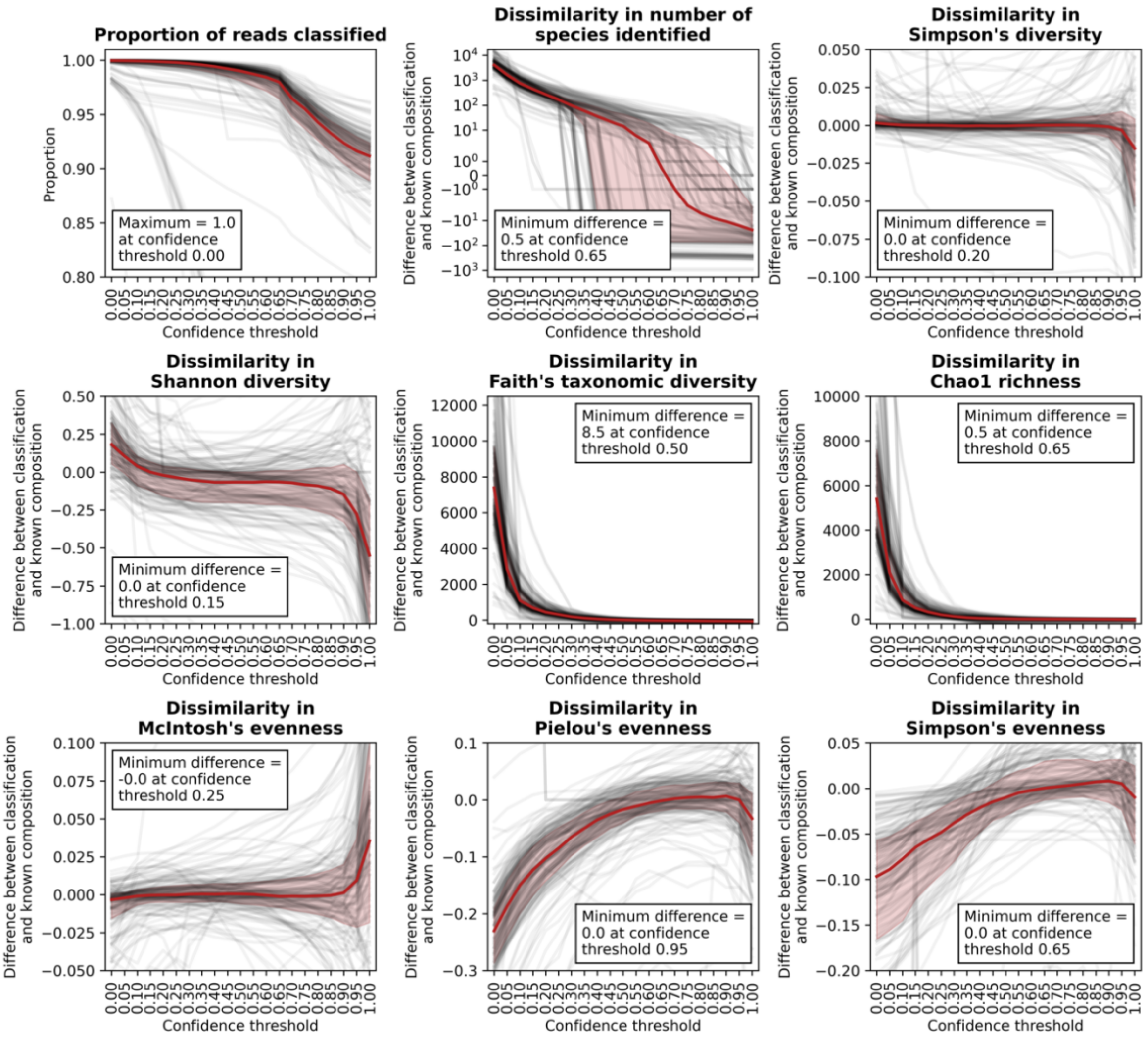


**Figure S4.** The proportion of reads classified and the dissimilarity in the number of species classified and alpha-diversity metrics for samples classified with Kraken2 with the NCBI RefSeq Complete V205 database at confidence thresholds between 0.00 and 1.00 at 0.05 intervals. All values shown here are the dissimilarity between the Kraken2 classifications and the known values, so as positive values show that the metric was higher in Kraken2 classifications than in the known composition, with negative values showing the opposite. Note here that we have used taxonomic rather than phylogenetic distance for Faith's taxonomic diversity (see Methods section). In each plot, black lines are for individual samples across different confidence thresholds and red lines and shaded areas show the median and upper and lower quartiles, respectively. Boxes with text indicate the optimal value (minimum difference between Kraken2-classification and known composition) and confidence threshold for each metric.

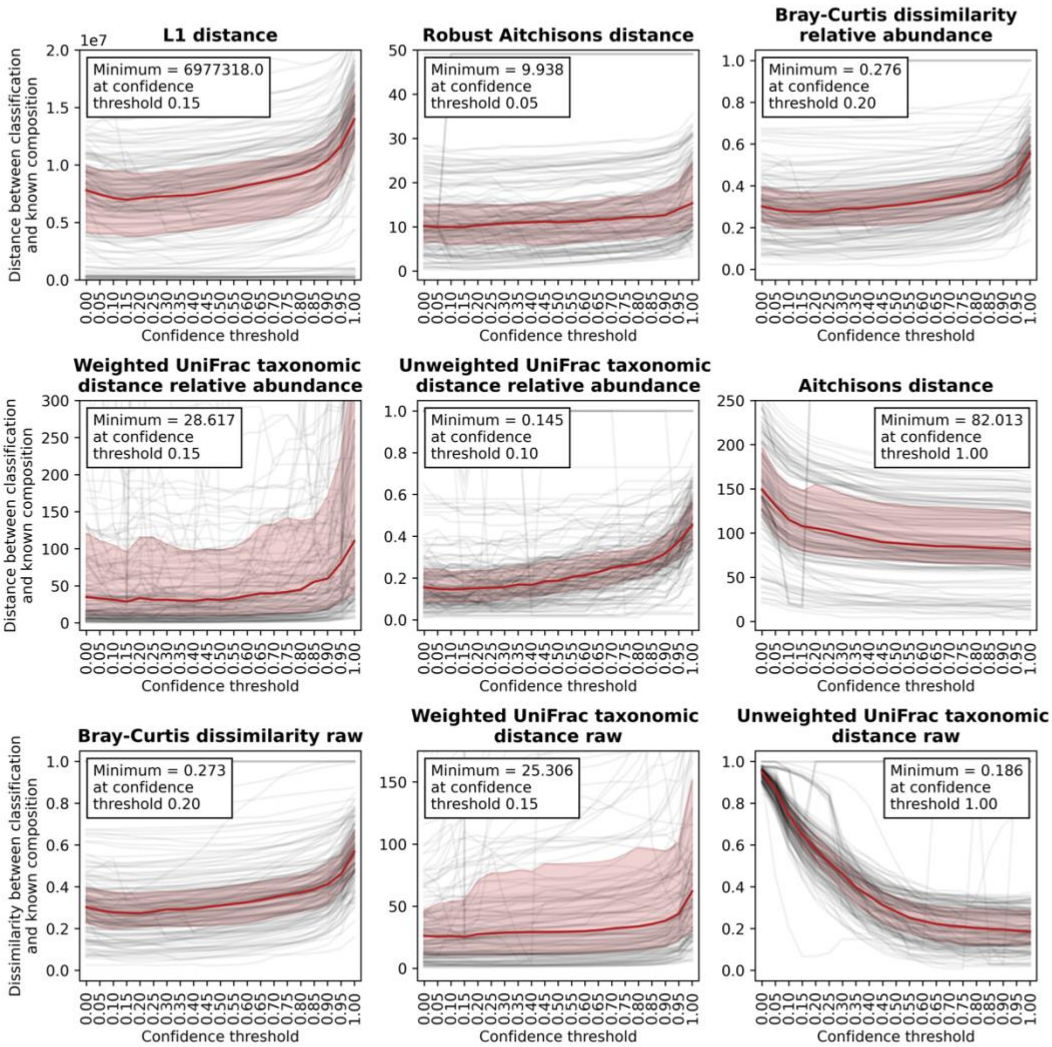


**Figure S5.** Beta-diversity metrics for samples classified with Kraken2 with the NCBI RefSeq Complete V205 database at confidence thresholds between 0.00 and 1.00 at 0.05 intervals. Note here that we have used taxonomic rather than phylogenetic distance for weighted and unweighted UniFrac distances (see Methods section). In each plot, black lines are for individual samples across different confidence thresholds and red lines and shaded areas show the median and upper and lower quartiles, respectively. Boxes with text indicate the optimal value (minimum distance) and confidence threshold for each metric.

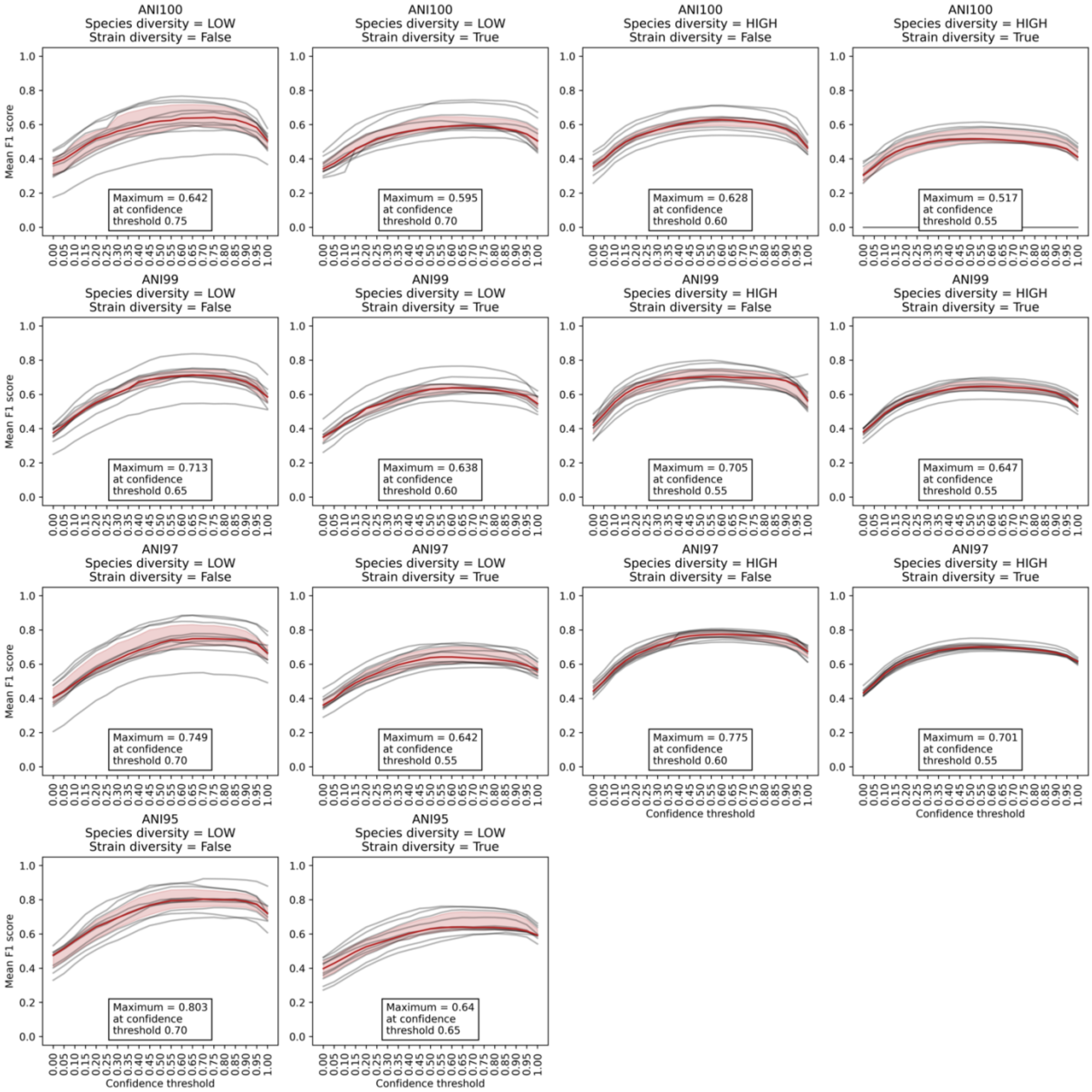


**Figure S6.** The effect of varying sample characteristics on the mean F1 score when Kraken2 is run with the NCBI RefSeq Complete V205 database at confidence thresholds between 0.00 and 1.00 at intervals of 0.05. Here, either the average nucleotide identity (ANI) between the genomes in the simulated samples and those in the databases is varied between 95% (ANI95) and 100% (ANI100), the species diversity is low (average of 100 species per sample) or high (average of 500 species per sample) or strain diversity is either present (up to 10 strains per species) or absent (one strain per species). In each plot, black lines are for individual samples across different confidence thresholds and red lines and shaded areas show the median and upper and lower quartiles, respectively. Boxes give the maximum mean F1 score and the confidence threshold at which this was obtained.

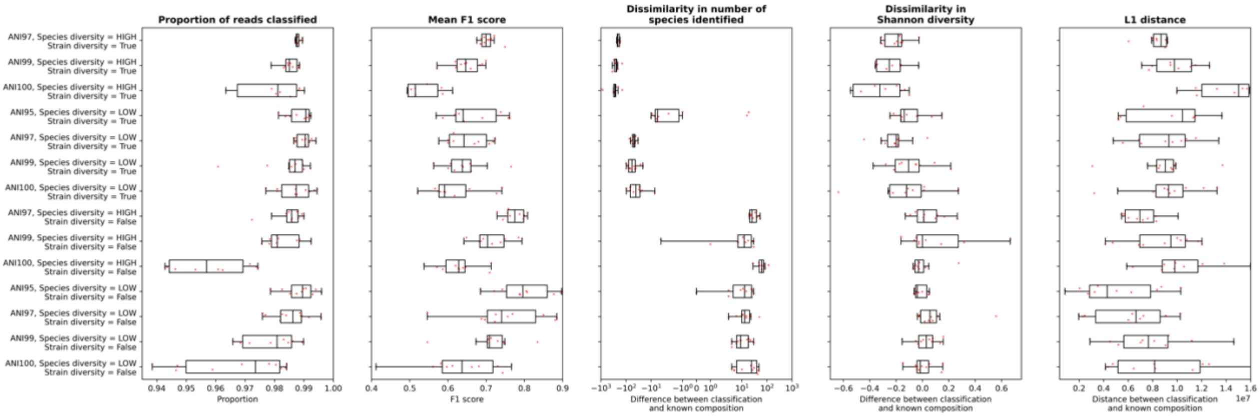


**Figure S7.** The effect of varying sample characteristics on the proportion of reads classified, mean F1 score, dissimilarity in number of species identified and Shannon diversity and L1 distance when Kraken2 is run with the NCBI RefSeq Complete V205 database at a confidence threshold of 0.60 (overall optimal mean F1 score). In each plot, each sample is shown as a single point with boxplots showing the median, upper and lower quartiles while whiskers show the range of the data (1.5 times the interquartile range). Here, either the average nucleotide identity (ANI) between the genomes in the simulated samples and those in the databases is varied between 95% (ANI95) and 100% (ANI100), the species diversity is low (average of 100 species per sample) or high (average of 500 species per sample) or strain diversity is either present (up to 10 strains per species) or absent (one strain per species).



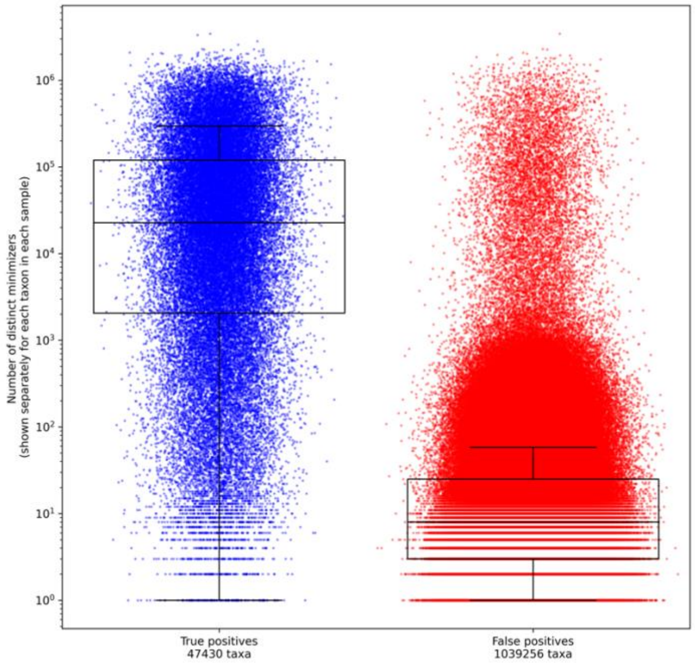


**Figure S8.** The number of distinct minimizers for true positive and false positive taxa for all simulated and mock samples run with Kraken2 and the NCBI RefSeq Complete V205 database (with no confidence threshold). Here, each point is for a single taxon in a single sample, with blue showing true positive taxa (taxa that were both classified by Kraken2 and found in the known sample composition) and red showing false positive taxa (taxa that were classified by Kraken2 but were not in the known sample composition). Boxplots show the median, upper and lower quartiles while whiskers show the range of the data (1.5 times the interquartile range).

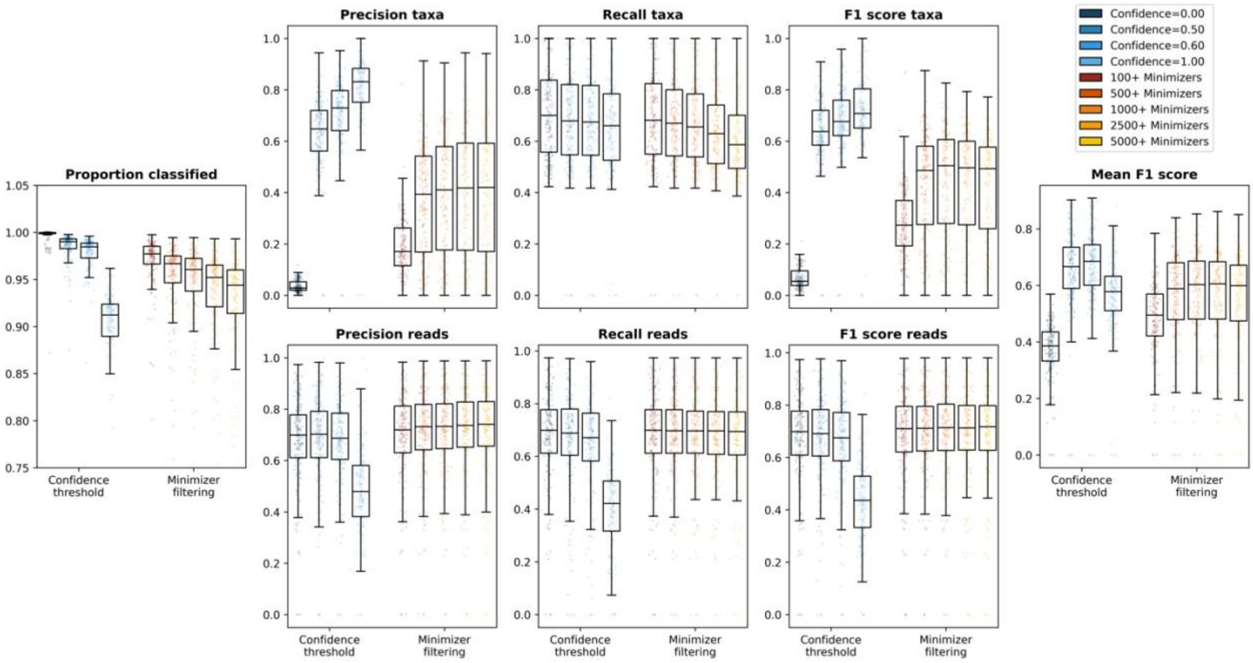


**Figure S9.** Comparison of using confidence thresholds or filtering taxa based on the number of distinct minimizers when simulated and mock samples were run with Kraken2 and the NCBI RefSeq Complete V205 database. Here, either a confidence threshold of 0.00, 0.50, 0.60 or 1.00 (blue colours) was used or taxa with below 100, 500, 1,000, 2,500 or 5,000 distinct minimizers (red colours) were removed from the classifications. In each plot, each sample is shown as a single point with boxplots showing the median, upper and lower quartiles while whiskers show the range of the data (1.5 times the interquartile range).

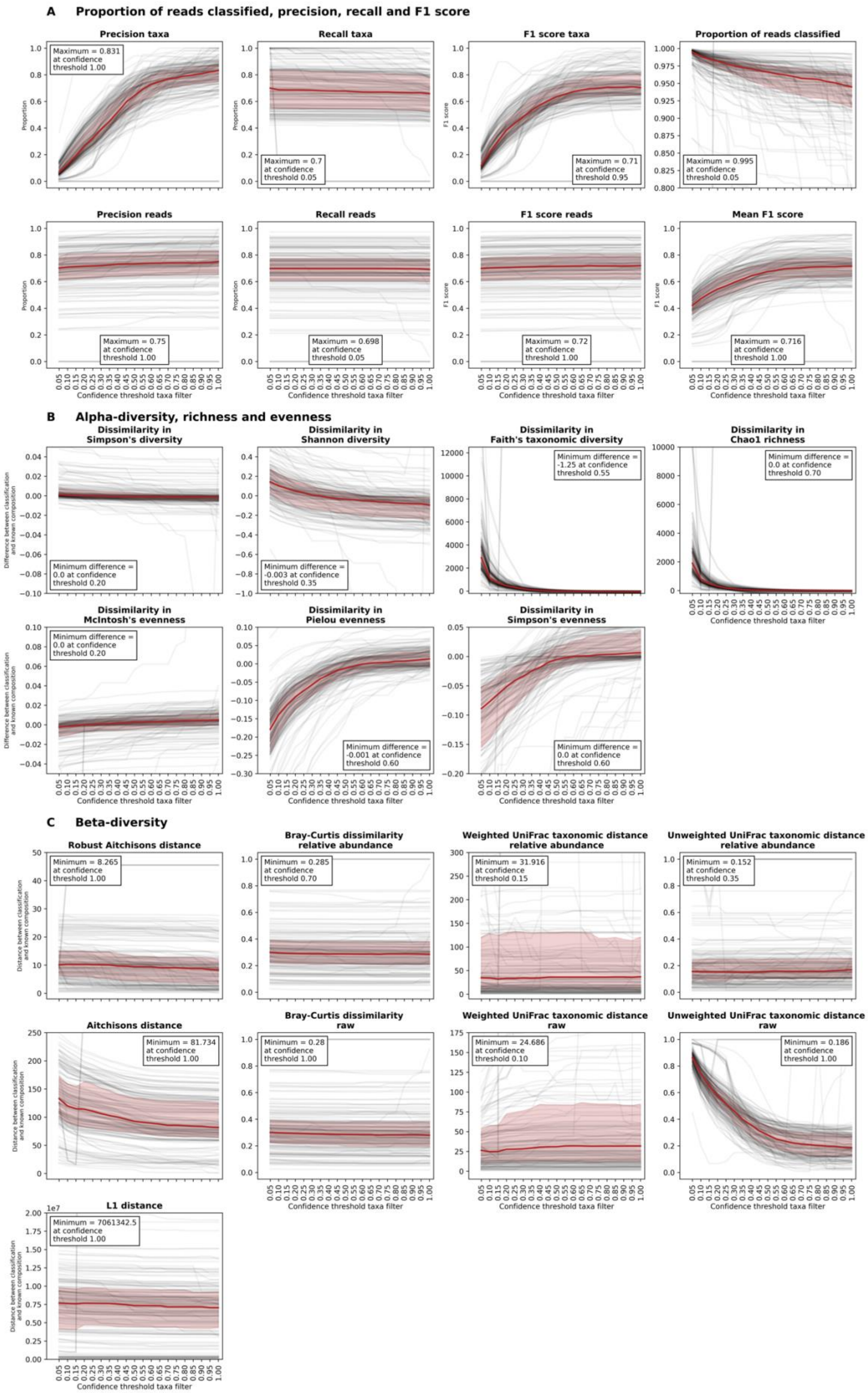
A Proportion of reads classified, precision, recall and F1 score
Precision taxa
Maximum = 0.831 at confidence threshold 1.00
Recall taxa
Maximum = 0.7 at confidence threshold 0.05
F1 score taxa
Maximum = 0.71 at confidence threshold 0.95
Proportion of reads classified
Maximum = 0.995 at confidence threshold 0.05
Precision reads
Maximum = 0.75 at confidence threshold 1.00
Recall reads
Maximum = 0.698 at confidence threshold 0.05
F1 score reads
Maximum = 0.72 at confidence threshold 1.00
Mean F1 score
Maximum = 0.716 at confidence threshold 1.00
Proportion
F1 score
Confidence threshold taxa filter
B Alpha-diversity, richness and evenness
Dissimilarity in Simpson's diversity
Minimum difference = 0.0 at confidence threshold 0.20
Dissimilarity in Shannon diversity
Minimum difference = -0.003 at confidence threshold 0.35
Dissimilarity in Faith's taxonomic diversity
Minimum difference = -1.25 at confidence threshold 0.55
Dissimilarity in Chao1 richness
Minimum difference = 0.0 at confidence threshold 0.70
Dissimilarity in McIntosh's evenness
Minimum difference = 0.0 at confidence threshold 0.20
Dissimilarity in Pielou evenness
Minimum difference = -0.001 at confidence threshold 0.60
Dissimilarity in Simpson's evenness
Minimum difference = 0.0 at confidence threshold 0.60
Difference between classification and known composition
C Beta-diversity
Robust Aitchisons distance
Minimum = 8.265 at confidence threshold 1.00
Bray-Curtis dissimilarity relative abundance
Minimum = 0.285 at confidence threshold 0.70
Weighted UniFrac taxonomic distance relative abundance
Minimum = 31.916 at confidence threshold 0.15
Unweighted UniFrac taxonomic distance relative abundance
Minimum = 0.152 at confidence threshold 0.35
Aitchisons distance
Minimum = 81.734 at confidence threshold 1.00
Bray-Curtis dissimilarity raw
Minimum = 0.28 at confidence threshold 1.00
Weighted UniFrac taxonomic distance raw
Minimum = 24.686 at confidence threshold 0.10
Unweighted UniFrac taxonomic distance raw
Minimum = 0.186 at confidence threshold 1.00
L1 distance
Minimum = 7061342.5 at confidence threshold 1.00
Distance between classification and known composition
Confidence threshold taxa filter

**Figure S10.** Performance metrics for samples run with Kraken2 against the full database built with NCBI RefSeq Complete V205. Rather than using the built-in Kraken2 confidence thresholds, here we have used the classifications given for no confidence threshold, but have filtered the reads so as only taxa present at the confidence thresholds of 0.05-1.00 (at intervals of 0.05) remain. In each plot, black lines are for individual samples across different confidence threshold filters and red lines and shaded areas show the median and upper and lower quartiles, respectively. (**A**) shows basic metrics including the proportion of reads classified as well as precision, recall and F1 score based on either taxa or reads. The mean F1 score is calculated as the mean of the F1 score based on taxa and the F1 score based on reads for each sample. (**B**) shows the dissimilarity in alpha-diversity, richness or evenness metrics between Kraken2-classified samples and the known composition for Simpson's diversity, Shannon diversity, Faith's taxonomic diversity, Chao1 richness, McIntosh's evenness, Pielou evenness and Simpson's evenness. Positive values show that the metric was higher in Kraken2 classifications than in the known composition, with negative values showing the opposite. (**C**) shows the distance or dissimilarity between Kraken2 classifications and the known composition for the L1 distance (also known as Manhattan distance), Robust Aitchison's distance, Aitchison's distance, Bray-Curtis dissimilarity (calculated with either raw or relative abundance values) and Weighted and Unweighted taxonomic UniFrac distance (both calculated with either raw or relative abundance values). Note here that we have used taxonomic rather than phylogenetic distance for both Faith's taxonomic diversity and weighted and unweighted UniFrac distances (see Methods section).

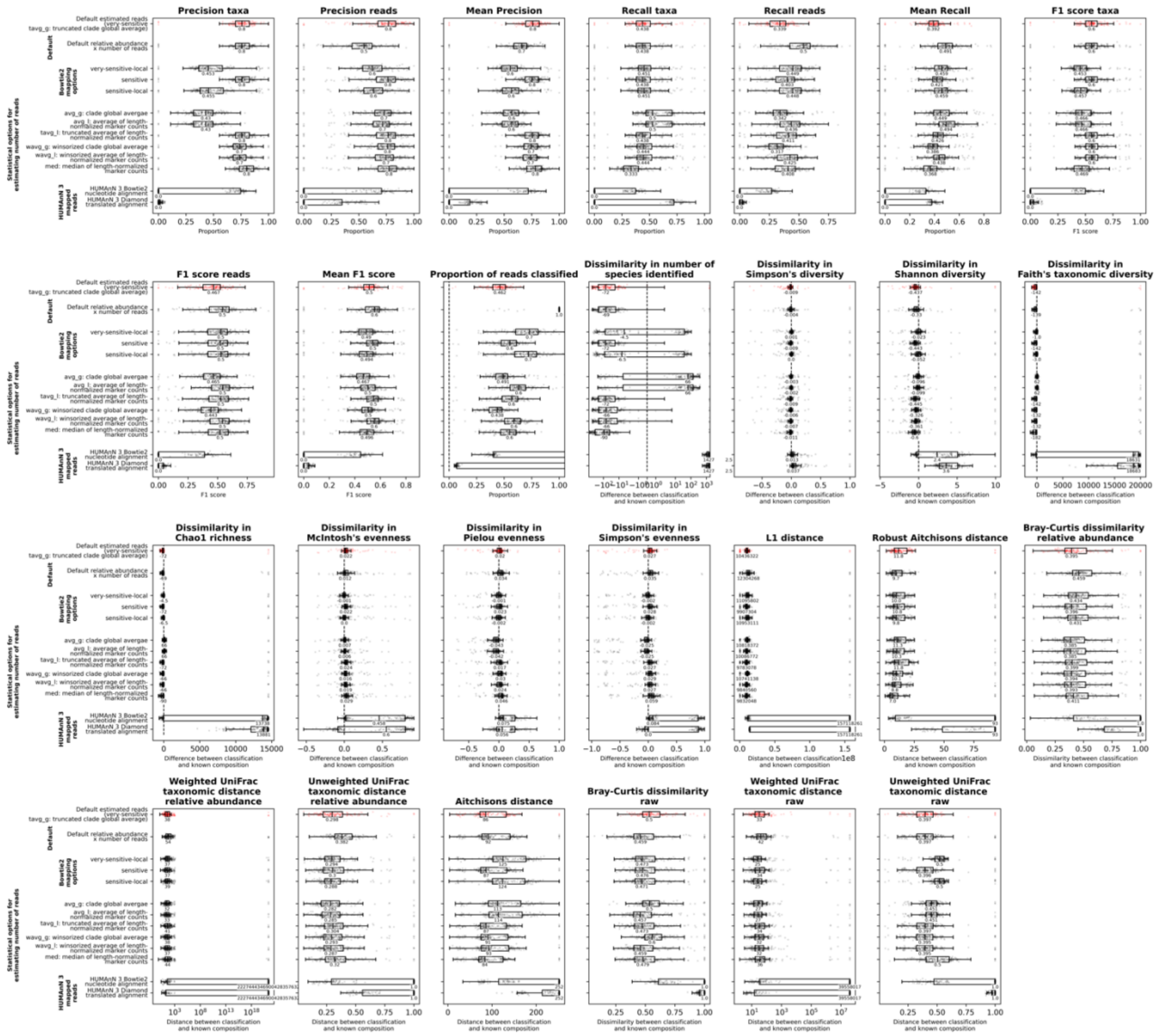


**Figure S11.** Calculated metrics for samples run using MetaPhlAn 3 with different parameters. In each plot, each sample is shown as a single point with boxplots showing the median, upper and lower quartiles while whiskers show the range of the data (1.5 times the interquartile range) and the default estimated number of reads is shown in red. This shows a comparison of the MetaPhlAn 3 default relative abundance multiplied by the number of reads in each sample (top group), using different Bowtie2 options for mapping reads to the marker gene databases (second group from top), using different statistical methods for estimating the number of reads contributed by each taxon/clade (second group from bottom) and using the reads that are mapped by different steps of the HUMAnN 3 pipeline (bottom group).

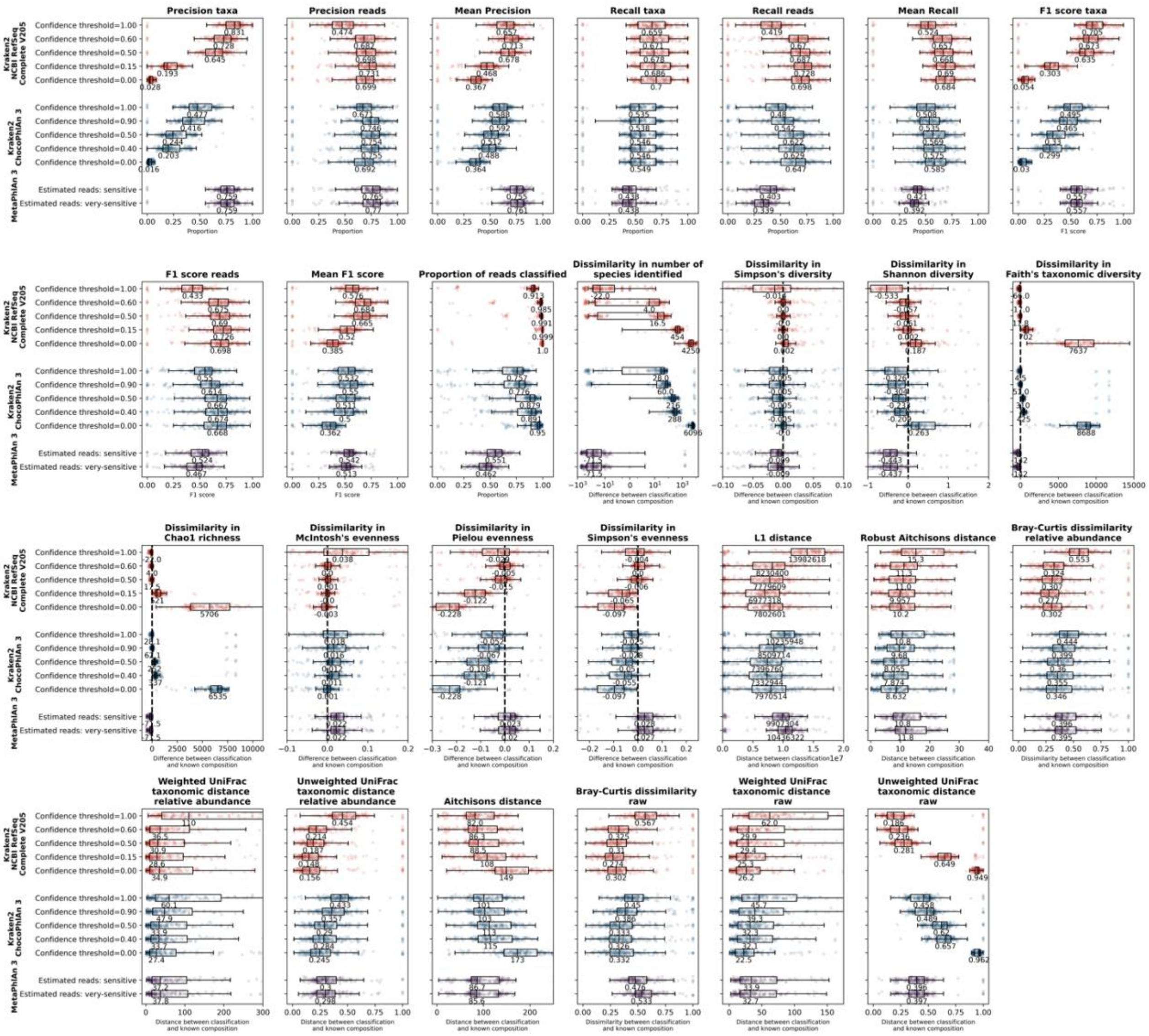


**Figure S12.** A comparison of the performance of MetaPhlAn 3 and Kraken2 with either the ChocoPhlAn 3 or the NCBI RefSeq Complete V205 databases, showing all metrics that were calculated. In each plot, each sample is shown as a single point with boxplots showing the median, upper and lower quartiles while whiskers show the range of the data (1.5 times the interquartile range). Kraken2 run with the NCBI RefSeq Complete V205 databases (top) is shown in red, Kraken2 run with the ChocoPhlAn 3 database (middle) is shown in blue and MetaPhlAn 3 (bottom) is shown in purple. For the two Kraken2 databases we have shown the results for running with confidence thresholds of 0.00, 0.50 and 1.00 in addition to the threshold at which the mean F1 score (0.60 for NCBI RefSeq Complete V205 and 0.90 for ChocoPhlAn 3) or L1 distance (0.15 for NCBI RefSeq Complete V205 and 0.40 for L1 distance) was optimised. For MetaPhlAn 3 we have shown the default estimated reads (using the very-sensitive Bowtie2 option) as well as the option where the L1 distance was optimised and the mean F1 score was almost optimised (the estimated reads with the sensitive Bowtie2 option).

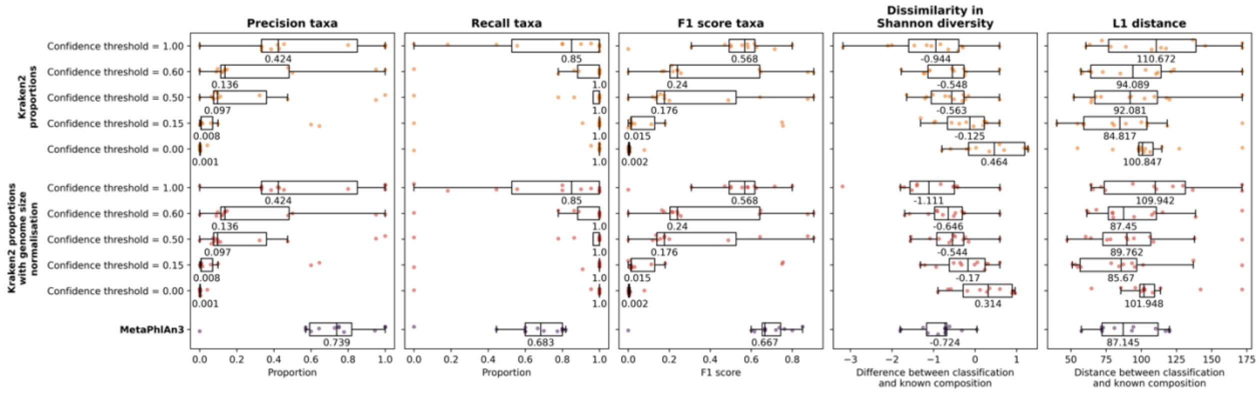


**Figure S13.** Comparison of MetaPhlAn 3 and Kraken2 with the NCBI RefSeq Complete V205 database run on mock samples with a known proportion of taxa, showing precision, recall and F1 score based on the taxa classified as well as dissimilarity in Shannon diversity and L1 distance between the classified and known compositions. Kraken2 results are shown both with and without genome size normalization, for confidence thresholds of 0.00, 0.15, 0.50, 0.60 and 1.00 (shown on the y-axis). In each plot, each sample is shown as a single point with boxplots showing the median, upper and lower quartiles while whiskers show the range of the data (1.5 times the interquartile range). Text underneath each box shows median values.

19

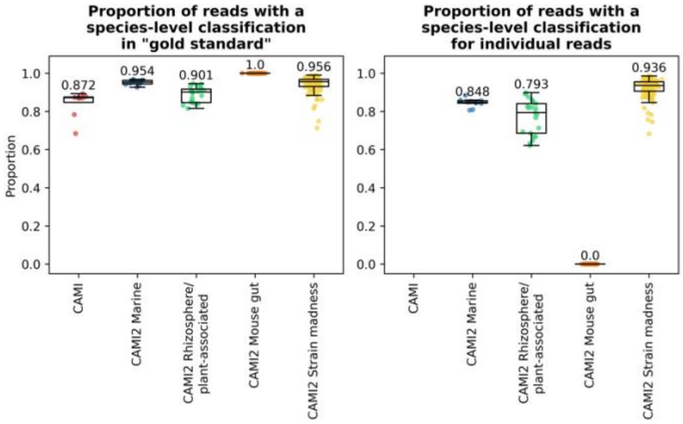


**Figure S14.** The proportion of reads with a species-level classification in the “gold standard” taxonomic profile for CAMI and CAMI2 samples of different types (left) or in the read-by-read classification for CAMI2 samples (right). Each sample is shown as a single point with boxplots showing the median, upper and lower quartiles while whiskers show the range of the data (1.5 times the interquartile range).

20

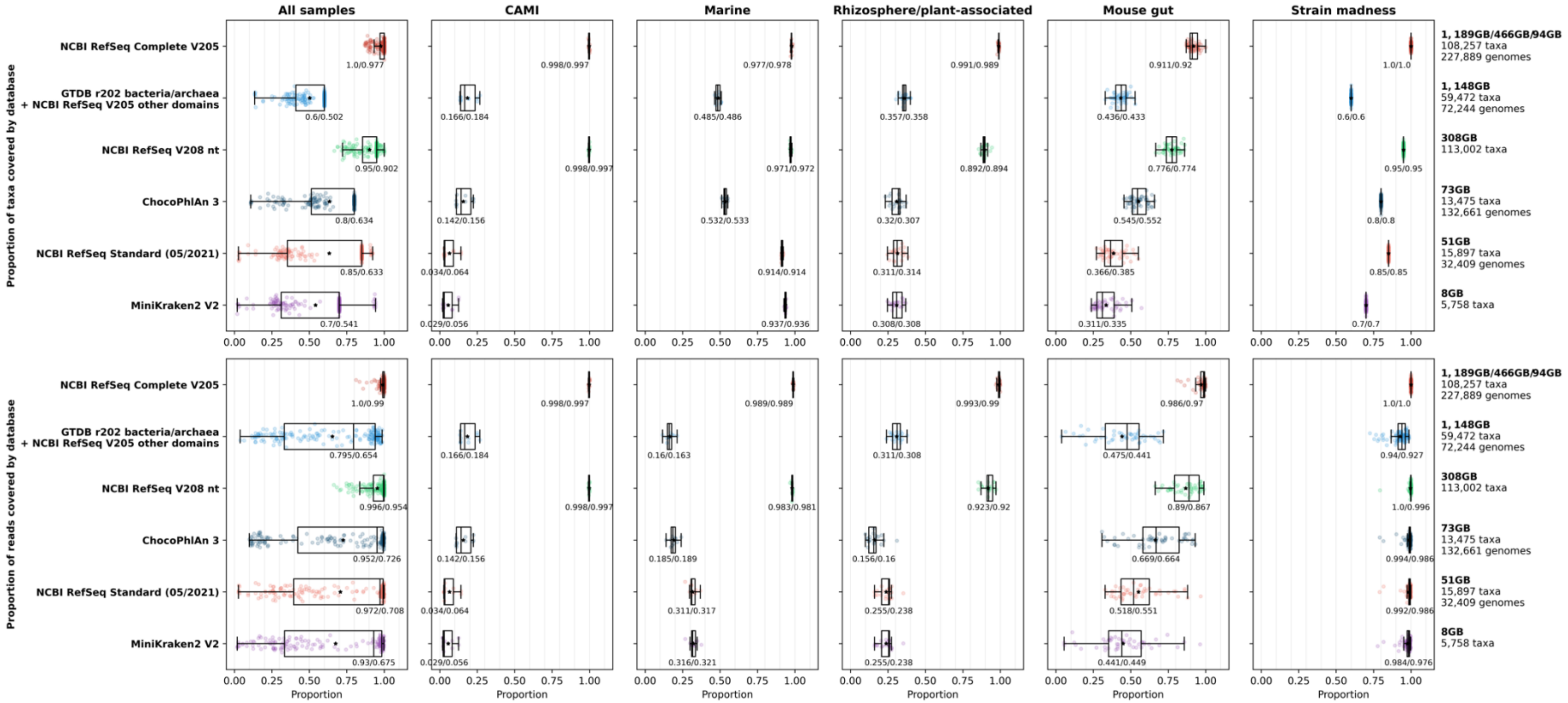


**Figure S15.** The proportion of taxa (top) and reads (bottom) within the "gold standard" taxonomic profiles of CAMI and CAMI2 samples that are covered by each database at the species level. All samples are shown together in the first column, and these are separated by sample type in the subsequent columns. Each sample is shown as a single point with boxplots showing the median, upper and lower quartiles while whiskers show the range of the data (1.5 times the interquartile range). Black stars indicate the mean for samples and numbers below the boxes show medians/means for all samples within each database. Databases are sorted by decreasing size from top to bottom. Full descriptions of the taxa contained within each database are given in the Methods section while a summary of the size, number of taxa and number of genomes (where this information is available) contained within each database is given on the right.

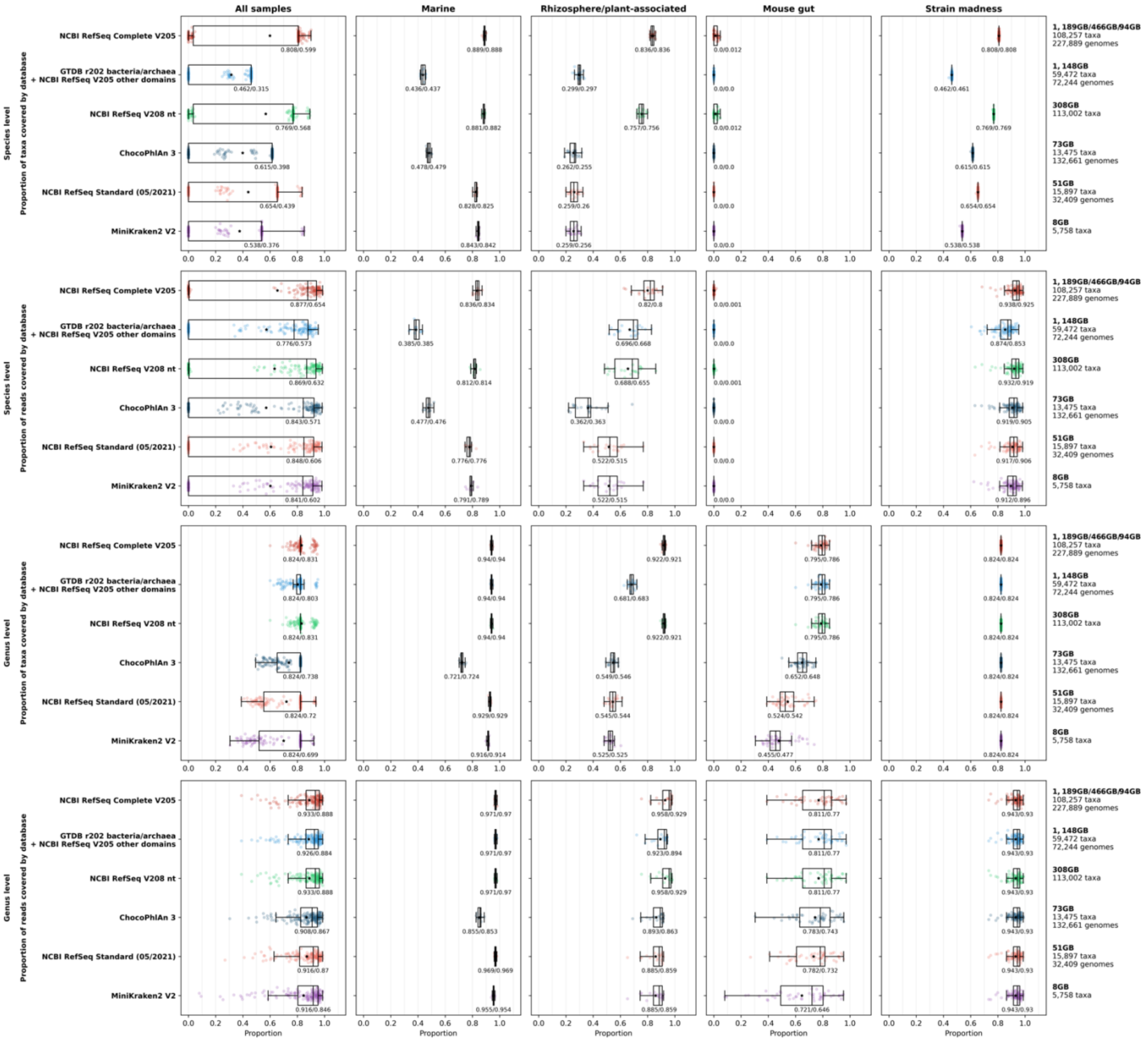
All samples
Marine
Rhizosphere/plant-associated
Mouse gut
Strain madness
Species level
Proportion of taxa covered by database
Species level
Proportion of reads covered by database
Genus level
Proportion of taxa covered by database
Genus level
Proportion of reads covered by database
NCBI RefSeq Complete V205
GTDB r202 bacteria/archaea + NCBI RefSeq V205 other domains
NCBI RefSeq V208 nt
ChocoPhlAn 3
NCBI RefSeq Standard (05/2021)
MiniKraken2 V2
1, 189GB/466GB/94GB
108,257 taxa
227,889 genomes
1, 148GB
59,472 taxa
72,244 genomes
308GB
113,002 taxa
73GB
13,475 taxa
132,661 genomes
51GB
15,897 taxa
32,409 genomes
8GB
5,758 taxa
Proportion

**Figure S16.** The proportion of taxa (first and third rows) and reads (second and fourth rows) within the read-by-read taxonomic profiles of CAMI2 samples that are covered by each database. This is shown for the species-level in the first two rows and the genus level in the third and fourth rows. All samples are shown together in the first column, and these are separated by sample type in the subsequent columns. Each sample is shown as a single point with boxplots showing the median, upper and lower quartiles while whiskers show the range of the data (1.5 times the interquartile range). Black stars indicate the mean for samples and numbers below the boxes show medians/means for all samples within each database. Databases are sorted by decreasing size from top to bottom. Full descriptions of the taxa contained within each database are given in the Methods section while a summary of the size, number of taxa and number of genomes (where this information is available) contained within each database is given on the right. Note that the proportions of taxa or reads covered at the species-level in mouse gut samples are close to zero because very few reads give a classification at the species rank.

22

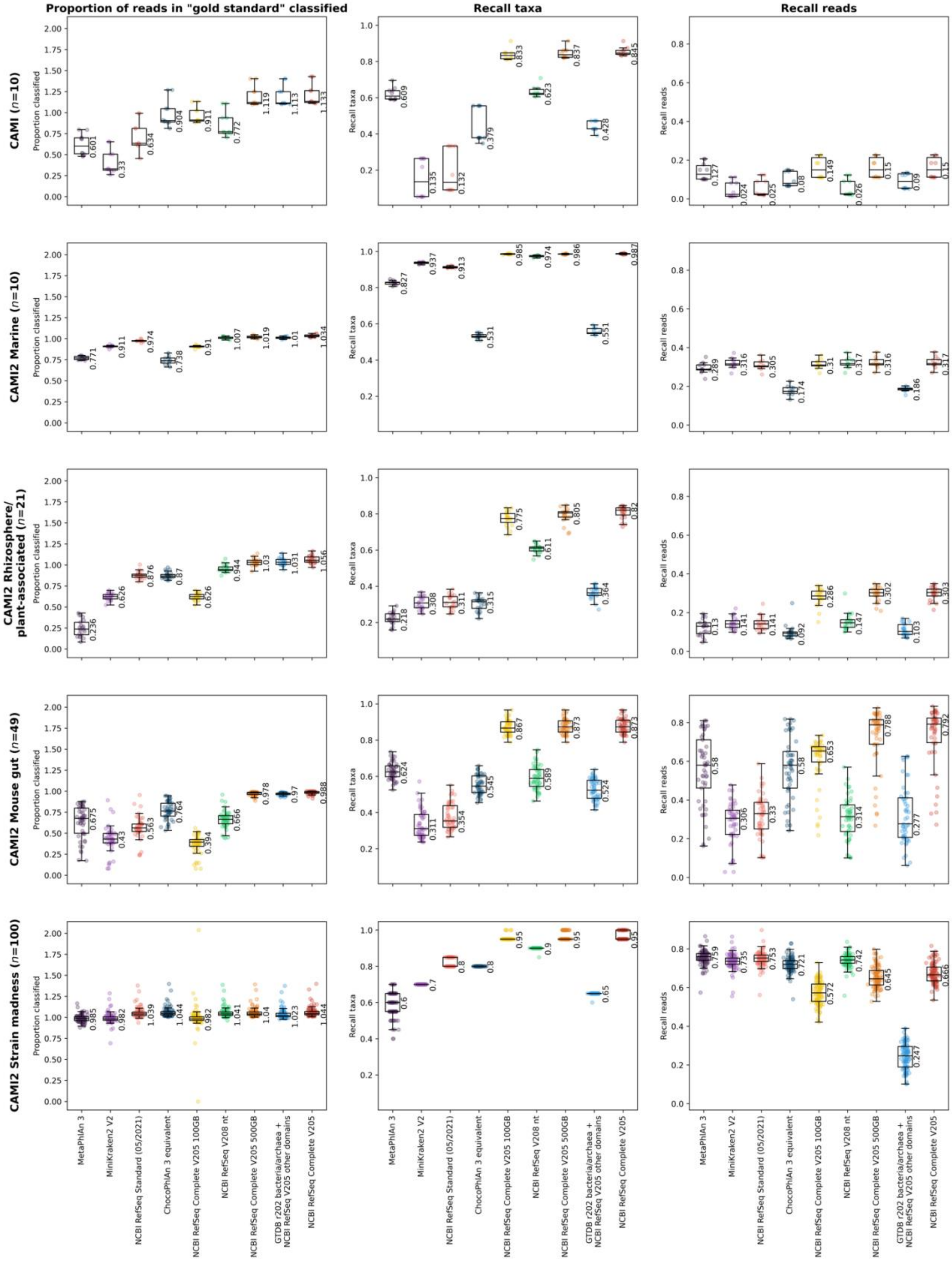


**Figure S17.** Proportion of reads in the "gold standard" taxonomic profiles classified (left), recall based on the taxa classified (centre) or recall based on the reads classified (right) for CAMI and CAMI2 samples run with MetaPhlAn 3 or Kraken2 with each of eight different databases. Note that the proportion of reads classified may be above 1 because the “gold standard” profiles do not include 100% of reads (Fig. S14). Rows show different sample types and the number of samples within this sample type. Each sample is shown as a single point with boxplots showing the median, upper and lower quartiles while whiskers show the range of the data (1.5 times the interquartile range) and values shown to the right of boxes are for the respective median. Databases are sorted by increasing size (left to right).

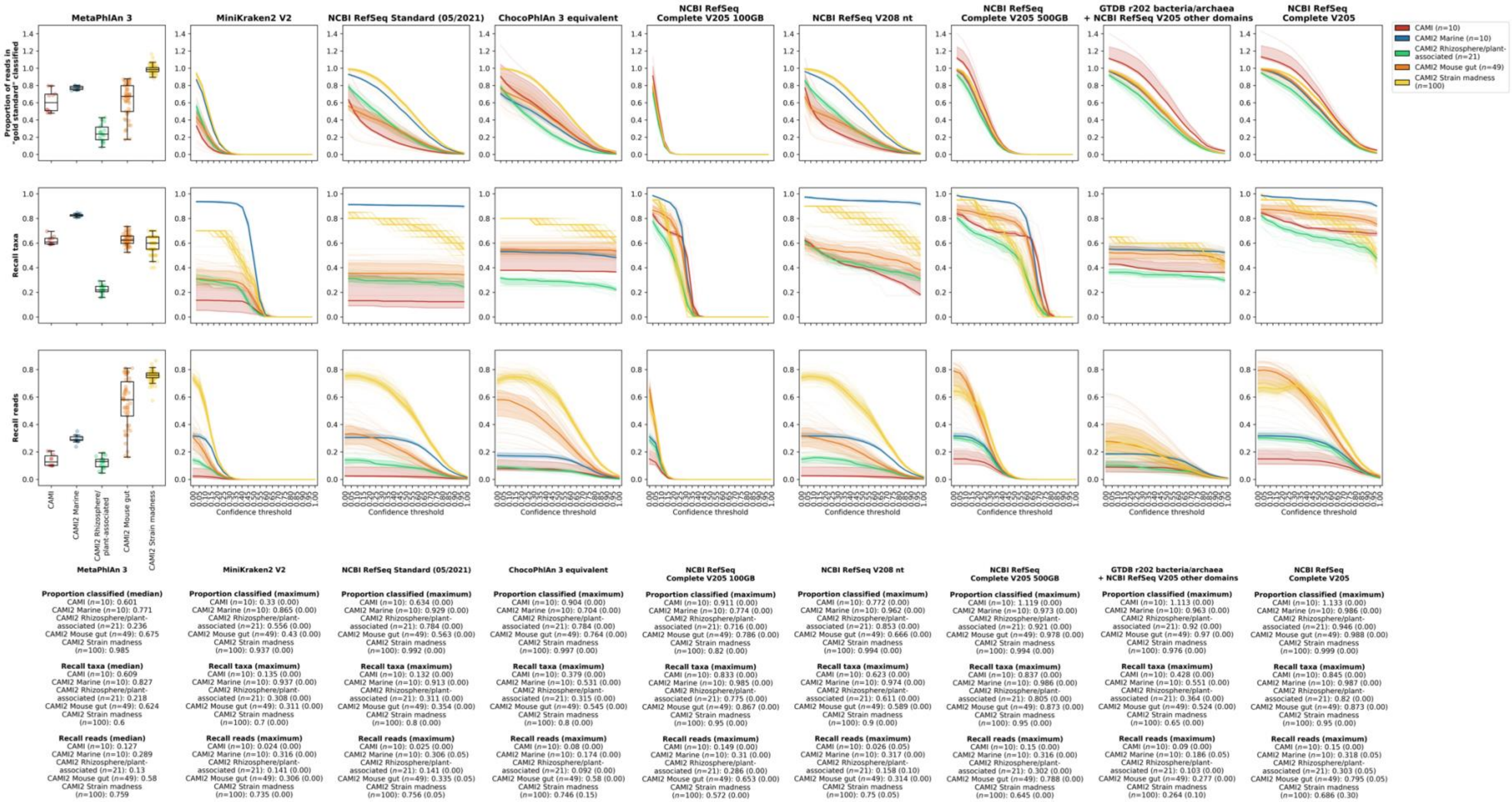


**Figure S18.** Proportion of reads in the "gold standard" taxonomic profiles classified (top row), recall based on the taxa classified (second row) or recall based on the reads classified (bottom row) for CAMI and CAMI2 samples run with MetaPhlAn 3 or Kraken2 with each of eight different databases (columns). For the MetaPhlAn 3 column, each sample is shown as a single point with boxplots showing the median, upper and lower quartiles while whiskers show the range of the data (1.5 times the interquartile range). For the Kraken2 columns, faint lines are for individual samples across different confidence thresholds (values between 0.00 and 1.00 at intervals of 0.05) while solid lines and shaded areas show the median and upper and lower quartiles, respectively, for each sample type. The median or maximum median value for each metric for each sample type is shown underneath the columns for each classification tool-database combination, for MetaPhlAn 3 or Kraken2, respectively, and for Kraken2 the number shown in brackets indicates the confidence threshold at which this maximum occurs. Databases are ordered by increasing size (left to right).

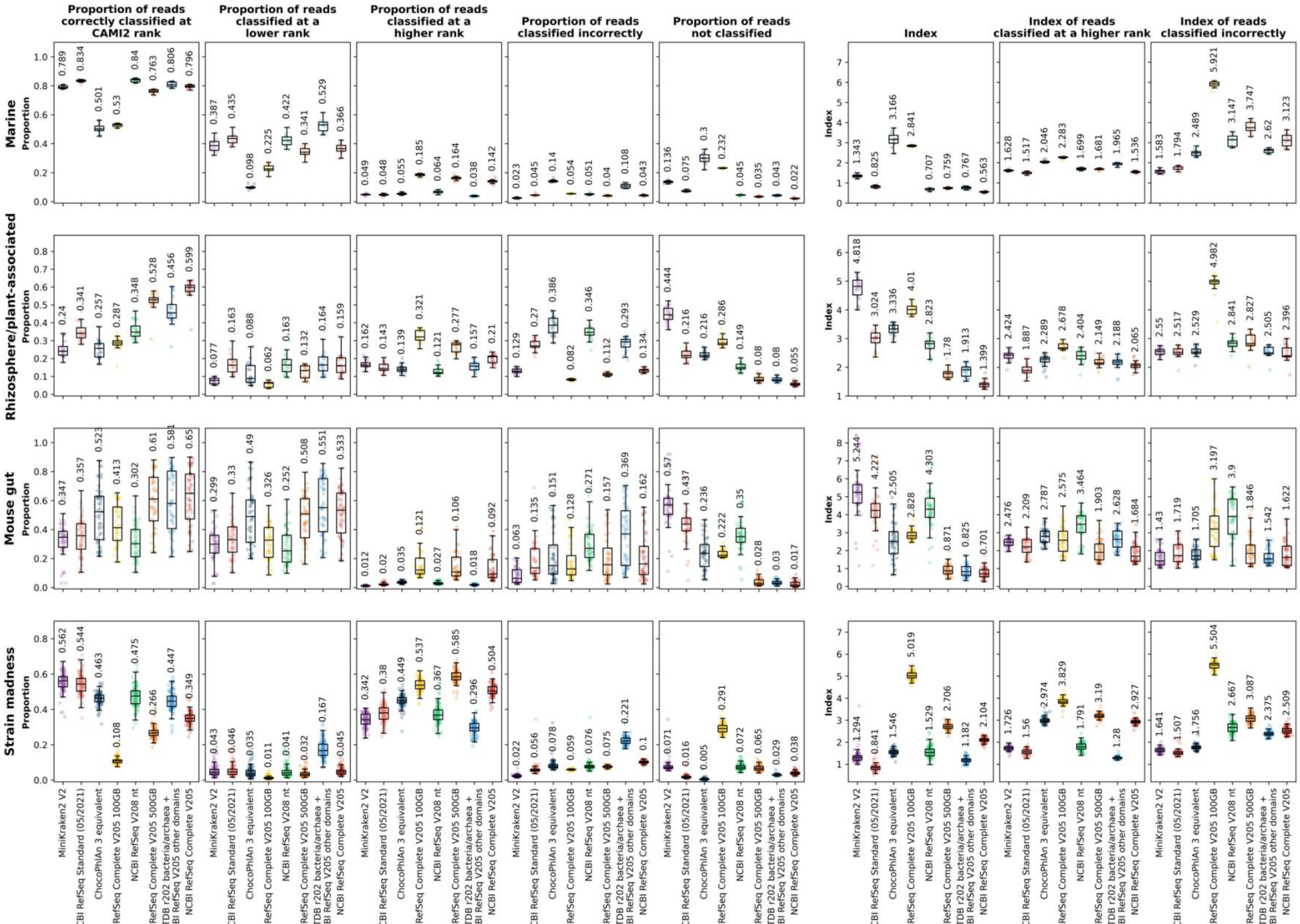

Proportion of reads correctly classified at CAMI2 rank
Proportion of reads classified at a lower rank
Proportion of reads classified at a higher rank
Proportion of reads classified incorrectly
Proportion of reads not classified
Index
Index of reads classified at a higher rank
Index of reads classified incorrectly
Marine
Rhizosphere/plant-associated
Mouse gut
Strain madness
Proportion
Index
MiniKraken2 V2
NCBI RefSeq Standard (05/2021)
ChocoPhlAn 3 equivalent
NCBI RefSeq Complete V205 100GB
NCBI RefSeq V208 nt
NCBI RefSeq Complete V205 500GB
GTDB r202 bacteria/archaea + NCBI RefSeq V205 other domains
NCBI RefSeq Complete V205

**Figure S19.** The proportion of reads correctly classified at the rank of the CAMI2 taxonomic classification (first column), the proportion of reads classified to a lower rank (second column), the proportion of reads classified to a higher rank (third column), the proportion of reads classified incorrectly (fourth column), the proportion of reads not classified (fifth column), the index for all reads (sixth column), the index of reads classified at a higher rank (seventh column) and the index of reads classified incorrectly (eighth column) for Kraken2 classifications of CAMI2 samples with all eight databases. Each sample is shown as a single point with boxplots showing the median, upper and lower quartiles while whiskers show the range of the data (1.5 times the interquartile range) for each database. Values shown above the boxes are for the respective median. Databases are sorted by increasing size from left to right.

### Supplementary results text

#### Effect of Kraken2 minimizer filtering on classification accuracy

Kraken2 includes an experimental feature that has been adapted from KrakenUniq [3] to give a count in the output of the number of minimizers and the number of distinct minimizers that a classification is based on. We initially looked at the overall number of distinct minimizers that were identified for true-positive *versus* false-positive taxa across all samples (Fig. S8), finding that the median number of minimizers found per taxon for true-positive taxa (*n*=22,703) was higher than for false-positive taxa (*n*=8). However, there were more false-positive (*n*=1,039,256) than true-positive (*n*=47,430) taxa identified in the samples and the maximum numbers of distinct minimizers identified per taxon was the same for true-positive and false-positive (*n*=3,441,048) taxa. It is likely for this reason that we didn't find filtering taxa based on the number of distinct minimizers to increase classification accuracy by much (Fig. S9), particularly in comparison with using a confidence threshold. Although there was an increase in median precision based on the number of taxa classified when we include all taxa (0.028) or only taxa with >100 (0.17) or >500 (0.394) distinct minimizers, this was relatively small compared with increasing the confidence threshold to 0.50 (0.648) or 0.60 (0.729). At the same time, the decrease in the median proportion of reads that were classified when we include all taxa (0.999) or only taxa with >100 (0.977) or >500 (0.967) distinct minimizers, was larger than when we increase the confidence threshold to 0.50 (0.99) or 0.60 (0.984). The decrease in recall based on the number of taxa and the changes in precision or recall based on the number of reads as the number of distinct minimizers was increased was comparable to that for increasing confidence threshold. Due to the described changes in precision based on the number of taxa classified and the proportion of reads classified, we did not investigate the impact of filtering taxa based on the number of distinct minimizers any further here.

#### Effect of filtering taxa based on confidence thresholds

We next investigated whether we could use the classifications that were obtained from running Kraken2 with the NCBI RefSeq Complete V205 database with no confidence threshold but removed any reads that belonged to taxa that were not present in the classified compositions at higher confidence thresholds (0.05-1.00). When using this confidence threshold filter, we found that the proportion of reads classified decreased steadily as the confidence threshold filter was increased, while the precision, recall and F1 scores based on taxa showed patterns that were similar to using the confidence threshold (Fig. S10 and Fig. 3 for the confidence threshold filter and confidence threshold, respectively). The precision, recall and F1 scores based on the number of reads did not change much with increasing confidence threshold filter, while the alpha- and beta-diversity metrics showed patterns that were similar to using the confidence threshold, but with less extreme values. Using the confidence threshold filter, we found that we could obtain a higher mean F1 score (0.705 at a confidence threshold filter of 1.00) than by using only the confidence threshold (0.684 at a confidence threshold of 0.60), although the L1 distance was also higher (7,410,771 at a confidence threshold filter of 1.00) than by using only the confidence threshold filter (6,977,318 at a confidence threshold of 0.15; Fig. S10 and Fig 2). Due to the higher optimal L1 distance obtained with the confidence threshold filter than with the confidence threshold, and the thought that filtering on a per-read basis is more useful than filtering on a per-taxon basis, we did not investigate the confidence threshold filter further.

#### Effect of varying MetaPhlAn 3 parameters on classification accuracy

While, to our knowledge, there has been no suggestion to vary parameters when running MetaPhlAn 3, as they were already comprehensively tested in the original study [4], we investigated the effect of varying several parameters within MetaPhlAn 3 and ran it in several different ways (described in the Methods section): (i) using the default parameters and either multiplying the resulting relative abundances by the number of reads in each sample or using the number of reads estimated by MetaPhlAn 3 with the rel_ab_w_read_stats option; (ii) using different options for Bowtie 2 mapping of reads to the marker gene database; (iii) using the different statistical options for estimating the number of reads in samples; and (iv) by running HUMAnN 3 and examining the number of reads mapped in both the Bowtie2 nucleotide alignment and Diamond-translated alignment steps. As above, for Kraken2, we compared the compositions given by MetaPhlAn 3 with the known compositions and calculated a range of different metrics based on these. Here we show only the proportion of reads classified, mean F1 score and the L1 distance (Fig. S12). The Simpson's diversity doesn't really vary based on these parameters because the number of taxa included is still the same (aside from with the different Bowtie2 mapping options, but this difference is negligible; Fig. S12). Note that the only instances where the proportion of reads classified is based on actual mapping of individual reads, rather than an estimate based on marker gene coverage, is in the HUMAnN 3 alignments, and that while we can compare the taxonomic composition of samples using these classifications, the majority would not be useful for downstream analyses that may wish to link taxonomy with other analyses carried out on individual reads.

### Impact of using default relative abundance vs estimated number of reads

We first compared the number of reads and found that multiplying the default relative abundance output of MetaPhlAn 3 by the number of reads within each sample had a higher mean F1 score than the number of reads belonging to each taxon estimated by MetaPhlAn 3 (default estimated reads; Fig. S11). Interestingly, even though we had set the unknown_estimation parameter in MetaPhlAn 3, it did not predict that a large proportion of the community by relative abundance was unknown, even though the median number of MetaPhlAn 3 estimated reads was less than 50% of the reads within samples. In this case, the precision and recall based on the number of taxa was the same for each so the difference in mean F1 score was based only on the precision and recall based on the number of reads, where the precision was higher for the MetaPhlAn 3 estimated reads and the recall was higher for the default relative abundance multiplied by the number of reads (Fig. S11). The L1 distance was lower for the MetaPhlAn 3 estimated reads.

### Options for Bowtie2 mapping of reads

We next investigated the impact of using different Bowtie2 options for mapping reads to the marker gene database on classification accuracy, where in each case we use the default MetaPhlAn 3 estimated reads. The proportion of reads that were classified was higher for the Bowtie2 options that used local mapping (Fig. S11), although it has previously been noted that using this option in MetaPhlAn 2 resulted in many spurious matches [5] and the sensitive option had both the highest mean F1 score and the lowest L1 distance anyway. This was due to the precision – reads and taxa – of both local options being much lower and the sensitive option had higher recall than the very-sensitive option, with similar (although slightly lower) precision (Fig. S11).

### Statistical options for estimating the number of reads

Another option that can be set within MetaPhlAn 3 is the statistical method used for estimating the number of reads contributed by each taxon. However, while we have presented these results here for comparison, we were unable to find a comprehensive description of what each of these options was actually doing. The method that gave the optimal value for each metric varied, however, the tavg_l (truncated average of length-normalized marker counts) method gave the joint highest mean

F1 score (with the wavg_l [winsorized average of length-normalized marker counts]) and the lowest L1 distance (Fig. S11). This suggests that the length-normalized methods perform better than the methods that don't account for length, but we recommend further investigation of these by any interested users as, while these options are given within MetaPhlAn 3, they are not what the authors recommend be used with MetaPhlAn 3.

#### HUMAnN 3 mapped reads

Many researchers that use MetaPhlAn 3 to taxonomically profile their metagenomic samples will also use HUMAnN 3 to functionally profile their samples. In HUMAnN 3, Bowtie2 is used to map reads from samples to ChocoPhlAn 3 pangenomes of the taxa that were identified by MetaPhlAn 3, that also include UniRef90 annotations. A further search is then performed against the database with the translated sequences using Diamond. We used the reads mapped in these steps to assess the number of reads that were classified using this pipeline, rather than the MetaPhlAn 3 estimated reads, as above (Fig. S11). While the metrics for the Bowtie2 aligned reads were similar to the MetaPhlAn 3 estimated reads, the proportion of reads classified and mean F1 scores were 0.05 and 0.04 lower and the L1 distance was 11,507,357 higher for the Bowtie2 aligned reads than the MetaPhlAn 3 estimated reads (10,595,954; Fig. S11). The proportion of reads classified and the mean F1 scores with the Diamond translated alignment was very low (0.06 and 0.04, respectively), while the L1 distance was very high (14,169,008).

#### MetaPhlAn 3 parameters to use

When choosing which MetaPhlAn 3 parameters to use, we have chosen not to consider the different statistical options for estimating the number of reads due to not finding clear guidance on how they were each calculated. Therefore, if the parameter with the highest mean F1 score (0.55) is chosen, then the best option is the default relative abundance multiplied by the number of reads, while if the lowest L1 distance (10,022,656) is chosen, then the best option is the sensitive Bowtie2 option (Fig. S12). We have, however, chosen the sensitive Bowtie2 option as well as the MetaPhlAn 3 estimated reads (default setting) for comparison with Kraken2 because: (a) the mean F1 score for the sensitive Bowtie2 option (0.54) is not much lower than the optimal; and (b) multiplying the default relative abundance by the number of reads gives an unrealistic number of reads classified by this method along with a much higher median L1 distance (12,444,744).

### Supplementary text references
